# Co-evolution of Large inverted repeats and G-quadruplex DNA in fungal mitochondria may facilitate mitogenome stability: the case of *Malassezia*

**DOI:** 10.1101/2023.02.07.527462

**Authors:** Anastasia C. Christinaki, Bart Theelen, Alkmini Zania, Selene Dall’ Acqua Coutinho, Javier F. Cabañes, Teun Boekhout, Vassili N. Kouvelis

## Abstract

Mitogenomes are essential due to their contribution to cell respiration. Recently they have also been implicated in fungal pathogenicity mechanisms. Members of the basidiomycetous yeast genus *Malassezia* are an important fungal component of the human skin microbiome, linked to various skin diseases, bloodstream infections, and increasingly implicated in gut diseases and certain cancers. In this study, the comparative analysis of *Malassezia* mitogenomes contributed to phylogenetic tree construction for all species. The mitogenomes presented significant size and gene order diversity which correlates to their phylogeny. Most importantly, they showed the inclusion of Large Inverted Repeats (LIRs) and G-quadruplex (G4) DNA elements, rendering *Malassezia* mitogenomes a valuable test case for elucidating the evolutionary mechanisms responsible for this genome diversity. Both LIRs and G4s coexist and convergently evolved to provide genome stability through recombination. This mechanism is common in chloroplasts but, hitherto, rarely found in mitogenomes.

---

Mitochondria are semi-autonomous organelles with their own genome, providing multiple vital processes, such as energy generation in the form of adenosine triphosphate (ATP) to the eukaryotic cell^1^. Fungi are lower eukaryotes found in every habitat and involved in all aspects of life and they act as decomposers, parasites, pathogens, or symbionts to other eukaryotic organism^2^. Fungal mitogenomes generally contain 14 protein-coding genes (*atp*6, *atp8, atp9, cob, cox*1-3, *nad*1-6 and *nad*4L), two ribosomal RNA (rRNA) genes (*rns* and *rnl*) and a variable number of transfer RNA genes (*trn*s), similarly to the mitogenomes of metazoa^1^. They may also encode other accessory genes like the ribosomal protein (*rps*3 or *var*1) and the RNA component for RNaseP (*rnp*B)^3,4^. Fungal mitochondrial genomes present a remarkable size variation ranging from 11,223 bp (*Hanseniaspora pseudoguilliermondii*)^5^ to 332,165 bp (*Golovinomyces cichoracearum*, NCBI Acc. No: NC_056148) and a variable gene order, which render them assets for studying fungal evolution^5–7^. This diversity of the mitogenomes is the result of the absence or presence of intergenic spacers, duplications, repeats, and insertions of plasmid components or other elements^4^. Conserved Large Inverted Repeats (LIR), although scarcely found, have been linked to mitogenome regulation^8^. Similarly, G-quadruplexes (G4) in mitogenomes are involved in the regulation of important biological processes, such us replication, altered gene expression, recombination, double-stranded break (DSB) formation, telomere regulation and genome instability^9,10^, as shown in the yeast *Saccharomyces cerevisiae*^8,11^.

Among the fungal species examined for their mt genome diversity, the basidiomycetous yeast *Malassezia* is an intriguing study case due to the mode of life of these yeasts^12^. It is the most abundant fungal genus on healthy human skin^13^ but may also cause various skin diseases such as atopic dermatitis, seborrheic dermatitis, dandruff, and pityriasis versicolor^14,15^. In recent years, *Malassezia* has increasingly been implicated in health and disease beyond the skin: as an underestimated cause of *Malassezia* bloodstream infections (BSIs) in immunocompromised patients and neonates^16^, as resident of the healthy human gut mycobiota^17^, but it has also been associated with Crohn’s disease^18^, colorectal cancer^19,20^, promoting pancreatic oncogenesis^21^, and is implicated in various respiratory diseases^22^. Several *Malassezia* species may also cause cutaneous pathologies in a variety of animals^12^. Direct sequencing approaches investigating various environmental sources, such as insects, nematodes, sponges, corals, soils, deep-sea vents and subglacial habitats, showed that this yeast may be ecologically more diverse than believed until now^12,23^.

*Malassezia* is lipid dependent with many fastidious species^24^, presenting challenges to obtain living cells from complex clinical and environmental sources which is needed to better understand the evolution of the genus and the mechanisms facilitating its role in various diseases and ecologies. With recent advancements in genomics, nuclear genomes are now available for all currently described species^24–27^ providing valuable resources for examining functional aspects and the evolutionary trajectory of this genus. However, only a limited number of studies have covered mitochondrial aspects and their contribution the evolution of the genus. Recently, *Malassezia sympodialis* and *Malassezia furfur* mitogenomes revealed LIRs^3,26,28^. In this study an effort was made to expand the analysis to all species of this genus. In detail, the first comprehensive analysis of the mitochondrial genomes of all 18 currently described *Malassezia* species plus two putative new ones provides insights into *Malassezia* mt genome organization and evolution is performed. Furthermore, it discusses the potential roles of LIRs, and G4s, raising the question whether these two features are functionally correlated and evolved convergently.

## Results

### General mitogenomic features reveal significant size variation and diversity

To date the mitochondrial genomes of nine *Malassezia* species are available in GenBank but only the ones of *M. furfur* and *M. sympodialis* have been described in detail^26,28^. In this study, we assembled and annotated the mitogenomes of the remaining 11 species, including a hitherto undescribed species from bat. In total, 28 mitogenomes were analyzed, representing all known *Malassezia* species including multiple strains, when available, for an in-depth pan-*Malassezia* comparative mitogenomic approach (Table 1).

**Table 1.**
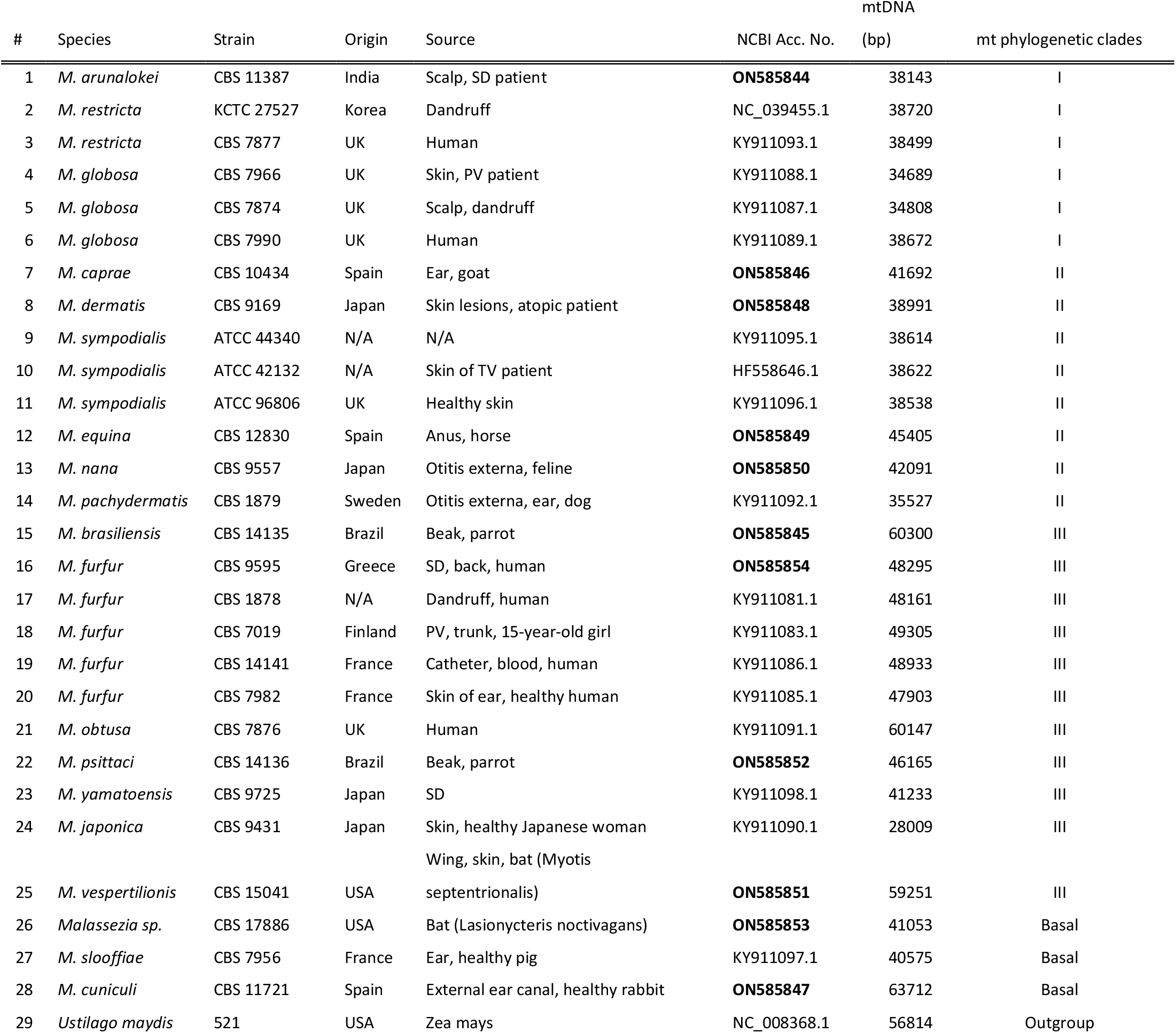
*Malassezia* species and strains used for the mitogenomic comparative analysis. Their origin, source is provided along with the NCBI Accession number and the size (in bp) of their mitogenome. The mitochondrial phylogenetic clade in which each species belong is also indicated. Bold lettering indicates the newly acquired mt genomes, sequenced and annotated in this work. NA: Non-Available, PV: pityriasis versicolor, SD: seborrheic dermatitis and TV: tinea versicolor.

Mitogenomes varied in size between 28,009 and 63,712 bp, with the one of *Malassezia japonica* being the smallest and the genome of *Malassezia cuniculi* the largest. This size variation mostly resulted from differences in number and size of introns, the presence of diverse LIRs and the size of intergenic regions (Suppl. Table S1). The number of introns varied between zero for *M. japonica* and two strains of *M. globosa*, and ten for hybrid *M. furfur* strain CBS 7019 with a total intron size ranging from zero to 9,172 bp (for *M. obtusa*). All introns were of group I type, located in the *rnl, cox1* and *cob* genes. Several introns hosted putative LAGLIDADG homing endonuclease genes (HEGs), with the exception of the first intron (ID subtype) in the *cob* gene of *M. pachydermatis* strain CBS 1879, which contained a GIY-YIG homing endonuclease gene (Suppl. Table S2).

The total number of nucleotides allocated to intergenic regions varied between 13,612 bp for *M. obtusa* and 47,344 bp for *M. cuniculi*. The GC-content was low as expected for mitochondrial genomes, approximately 30 %. The mitogenomes for all analyzed strains contained the expected standard 14 protein-coding genes, two rRNA genes, between 22 and 32 tRNAs, and the protein coding *rps*3 gene, with *M. slooffiae* and *M. cuniculi* containing two copies of this gene. *M. slooffiae* uniquely possessed the gene for the RNA subunit of mitochondrial RNase (*rnp*B). Most *Malassezia* mitogenomes contained a LIR with a variable number of duplicated genes (Suppl. Table S1).

For some species, mitogenomes for multiple strains were available, reanalyzed, and assessed, showing a variable level of intraspecies mitogenomic differences mostly resulting from the diversity of their intergenic regions and LIRs, and their intron abundance. For example, of the three *M. globosa* strains, the mitogenomes of the CBS 7966 and CBS 7874 do not possess any introns, but CBS 7991 contains three introns.

Up to now an intraspecies mitogenome variability was previously analyzed for 20 *M. furfur* strains^3^. The authors reported a mitogenome size variability ranging from 45,715 to 49,317 bp, mostly resulting from variation in intron abundance and variable intergenic regions^3^. In the present study the mitogenome of CBS 9595 was selected and analyzed, as this turned out to be an important parental reference strain in a study focusing on *M. furfur* hybridization events^28^.

### Mitochondrial phylogeny

Phylogenetic relationships were determined considering the 15 mt protein coding genes, with *Ustilago maydis* used as outgroup (Figure 1). Three main phylogenetic clades could be determined: clade I, containing *M. arunalokei, M. restricta*, and *M, globosa*; clade II, with *M. caprae, M. dermatis, M. sympodialis, M. equina, M. nana*, and *M. pachydermatis*; clade III, containing *M. brasiliensis, M. furfur, M. obtusa, M. psittaci, M. yamatoensis, M. japonica*, and *M. vespertilionis*; and finally, the basally positioned species *M. slooffiae* and *M. cuniculi*, together with strain CBS17886 which represents a putative new species. In current mitogenomic phylogeny, *M. furfur* CBS 9595 was positioned slightly deviating from the other four *M. furfur* strains and was found to be closer related to *M. brasiliensis*. Based on the concatenated 15 protein coding genes (PCGs), sequence similarity between CBS 9595 and the other four *M. furfur* strains was 98.3 - 98.4% compared to 99.7-99.9% among the four closer related strains, corroborating previous indications that *M. furfur* may comprise two closely related species^28^ (Suppl. Table S3).

**Figure 1.**
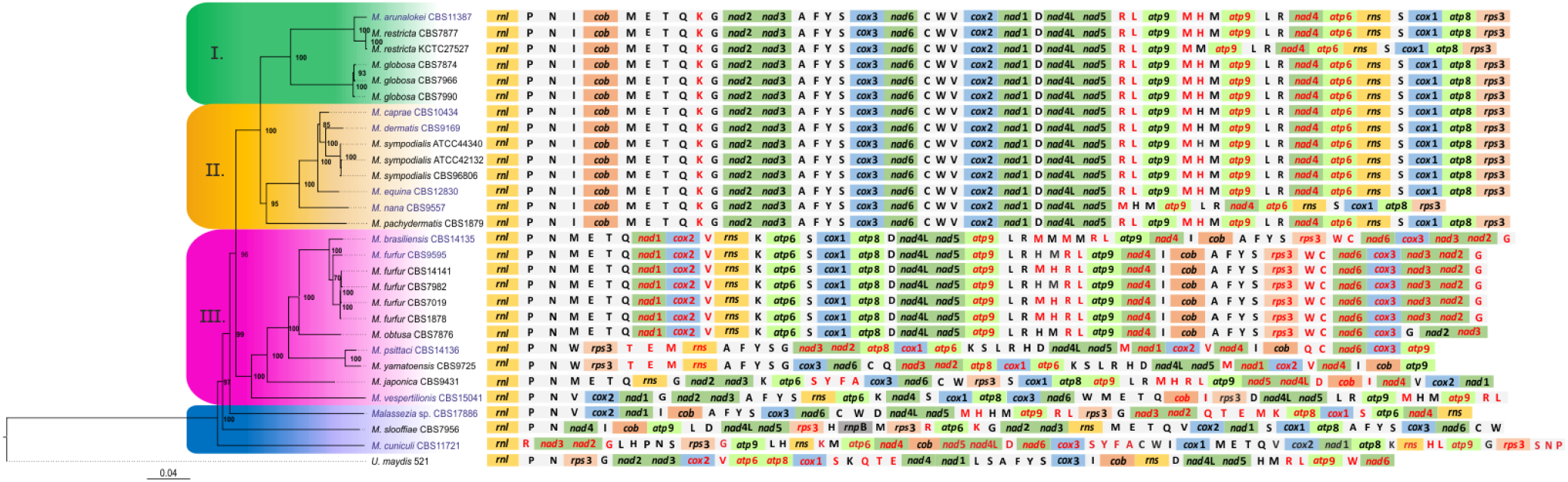
Phylogeny and synteny of the mitogenome of 28 *Malassezia* strains representing all currently known species. The phylogenetic tree was produced using Bayesian Inference based on the concatenated amino acid matrix of the 15 mitochondrial PCGs. Numbers at the nodes of the tree indicate posterior probability (PP) values. The species *Ustilago maydis* strain 521 has been used as an outgroup. Shading of phylogenetic clusters is shown as green, yellow, purple, and blue for clades I, II, III and the basal species, respectively. Blue text indicates the taxa whose mt genome is presented for the first time in this study. Additionally, synteny is provided in all cases starting with *rnl* gene. The genes which are located on the complementary reverse strand are marked with red color. Amino acid symbols represent the corresponding tRNA gene (e.g., P for *trn*P). Genes referring to subunits of NADH complex, apocytochrome b, cytochrome c oxidase complex and ATP synthase complex are shown in green, orange, blue, light green respectively.

### Mitochondrial synteny analyses suggest evolution towards fixated genome organization

Mitogenomic gene shuffling between species was observed, and only two groups of species presented a similar syntenic configuration (Figure 1). Species of phylogenetic clades I and II, except for *M. nana*, showed identical gene order indicating late evolutionary acquired genome conservation. Some LIR variation existed for *M. nana* and *M. restricta* strain KCTC 27527 (see next section). The other group with identical synteny consisted of the closely related species *M. brasiliensis, M. furfur*, and *M. obtusa* from phylogenetic group III. One deviation in genome organization could, however, be observed in *M. obtusa*, for which the *trn*G-*nad*2-*nad*3 gene cluster was positioned in an inverted orientation compared to *M. brasiliensis* and *M. furfur*. The remaining seven species retained unique genome organizations with only partially shared synteny. Among all *Malassezia* species, six syntenic blocks could be distinguished: *trn*M-*trn*E-*trn*T, *nad*2-*nad*3, *cox*3-*nad*6, *trn*V-*cox*2-*nad*1, *trn*D-*nad*4L-*nad*5, and *trn*A-*trn*F-*trn*Y-*trn*S (Figure 2, i-vi). If *M. cuniculi* is excluded, three more syntenic blocks could be identified (i.e., *rnl*-*trn*P-*trn*N, *trn*I-*cob* and *trn*S-*cox*1-*atp*8) (Figure 2, vii-ix). Syntenic diversity was illustrated by the transposition and inversion of gene blocks and single genes among species. These events most probably occurred among the earlier divergent species, with *M. cuniculi* mt genome containing the most differing genome organization. As phylogenetically basal species possessed a less organized gene order among them compared to the more evolved clusters, an evolutionary drive towards fixated genome organization seems to exist.

**Figure 2.**
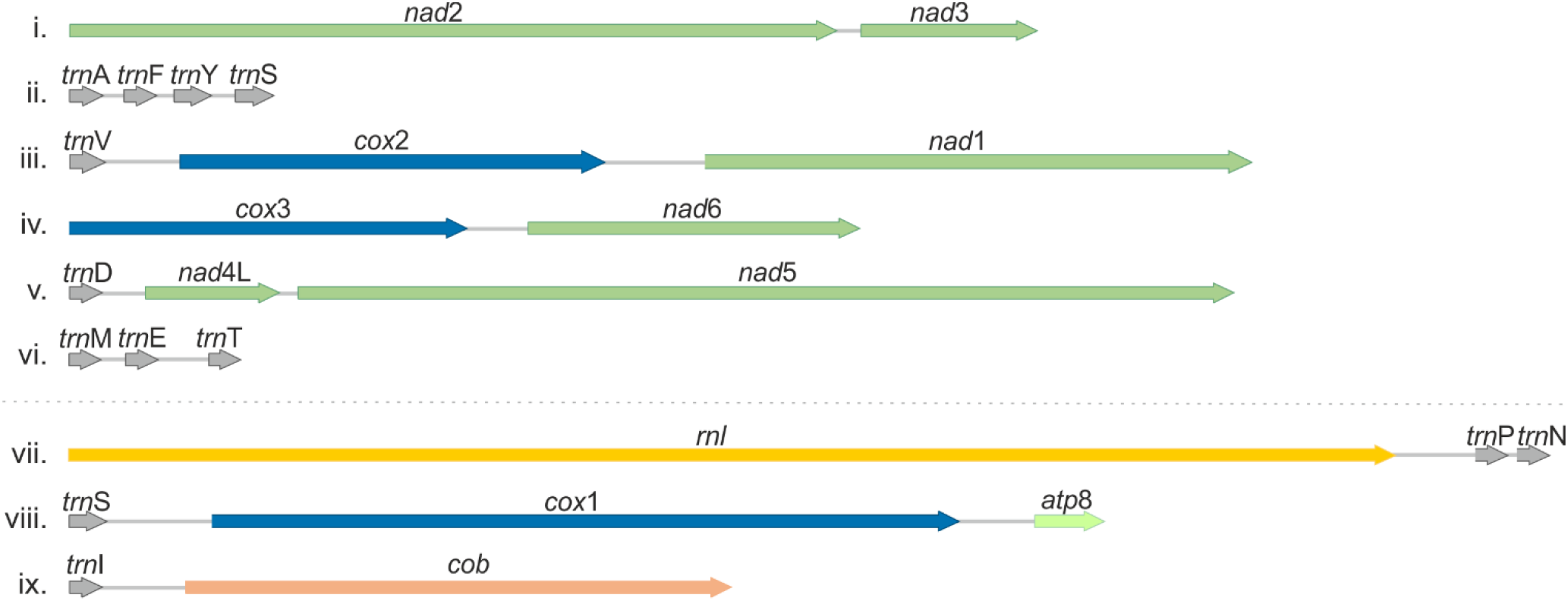
The main syntenic units of *Malassezia* mitogenomes. The first six units can be found in all *Malassezia* spp. (i.e., i-vi) while the last three in all except *M. cuniculi* (i.e., vii-ix). Arrows indicate gene transcription direction.

### Long Inverted Repeats (LIRs)contribute to the conserved synteny of the more recently evolved phylogenetic clades

Nearly all *Malassezia* species contained a LIR varying in size between 2,098 bp for *M. japonica* and 10,698 bp for *M. brasiliensis* (Suppl. Figure S1). Variation in LIR assembly resulted from the presence or absence of *trn* genes in the repeat unit and in the intra-IR region, the orientation or combinations of genes, and finally from variation in the intergenic regions (Figure 3).

**Figure 3.**
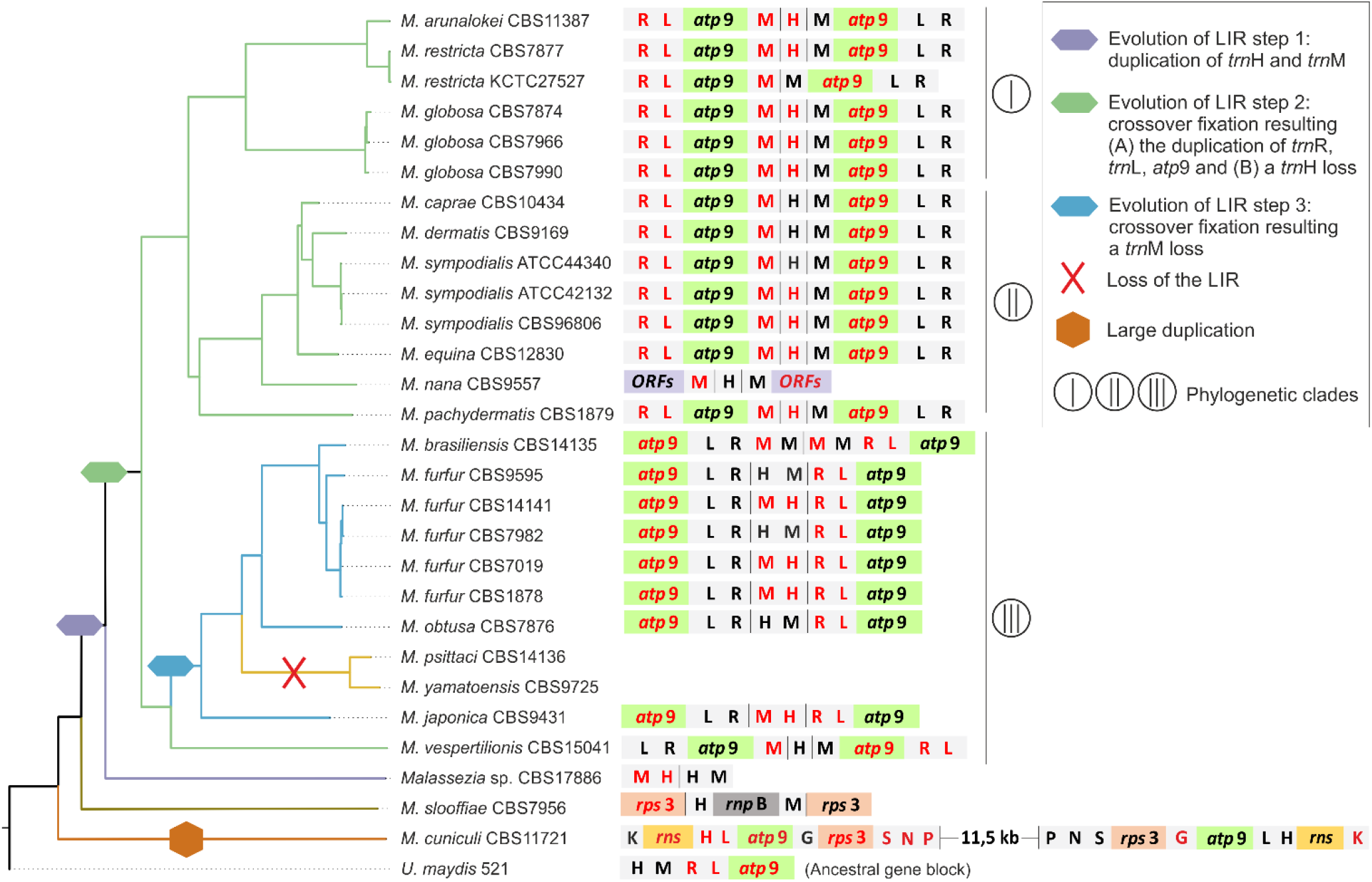
Schematic representation of *Malassezia* LIR evolution in relation to phylogeny. LIRs synteny is presented next to the respective phylogenetic clade. The intra-IR region can be seen in between black vertical lines. The genes located on the complementary reverse strand are indicated in red coloring and amino acid symbols represent the corresponding tRNA gene (e.g., H for *trn*H). The main evolutionary steps are shown as different colored symbols of the nodes and clades of the tree. Latin numbers in circles indicate the three different phylogenetic clades. The ancestral gene block of *Ustilago maydis* is provided.

The early diverged mt genomes of *M. cuniculi* and *M. slooffiae* evolved independently, resulting in unique synteny and LIR composition. In *M. cuniculi* a duplicated region composed by *trn*P-*trn*N-*trn*S-*rps3*-*trn*G-*atp*9-*trn*L-*trn*H-*rns*-*trn*K was present with the second copy positioned in an inverted orientation and separated by a long genomic stretch containing *rnl*-*trn*R-*nad*3-*nad*2-*trn*G-*trn*L-*trn*H (Figure 3). Thus, it should probably be disqualified as a LIR but attributed to a simple inverted duplication. This duplication may have been the result of a recombination event and it facilitated the existence of duplicated *rps*3 and *rns* gene copies. In *M. slooffiae* the repeat unit contained the gene *rps*3 with genes *trn*H, *rnp*B, *trn*M located in the intra-IR region. The protein coding gene *rnp*B is only present in the mitogenome of *M. slooffiae* and it seems that this gene was introduced with the insertion of the LIR.

In CBS17886, representing a yet undescribed species *Malassezia* species, a 3.7 kb fragment including the genes *trn*M and *trn*H was observed comprising the first evolutionary step for the formation of laterally evolved LIRs in all other *Malassezia* species. Genes included in *Malassezia*’s LIRs were also found in the *U. maydis* mitogenome as an ancestral gene block composed by *trn*H-*trn*M-*trn*R(inverted)-*trn*L(inverted)-*atp*9(inverted). Phylogenetic clades I and II differed from clade III in their LIR and intra-IR content. In clades I and II, *trnM-atp*9(inverted)-*trnL*-*trnR* constituted the LIR and the intra-IR included the *trnH* gene. In clade III (with the exception of *M. vespertilionis*), the LIR included only *trnR*(inverted)*-trnL*(inverted)*-atp*9, and the intra-IR consisted of *trnH*-*trnM*. Two species of clade III, *M. yamatoensis* and *M. psittaci*, lacked the LIR entirely, which may have been lost in a more recent evolutionary event, as both species clustered together in a separate clade (Figure 3).

In *M. furfur* and *M. sympodialis*, intraspecies LIR variation was also detected and mainly resulting from different orientations of the intra-IR fragment. Interestingly, one of the *M. restricta* strains lacked the intra-IR region.

### G-quadruplex (G4) motifs in *Malassezia* mitogenomes are predominantly distributed in the LIRs

Potential G4 formation was identified in all examined mitogenomes, varying from one in *M. yamatoensis* to 37 in *M. obtusa*, without significant correlation with the mitogenome size and the GC content (Suppl. Table S4, Suppl. File S1). G4s are mainly distributed within the LIRs, with few exceptions. *M. globosa* was the only species without any G4s in the LIR (Figure 4). In the *M. yamatoensis* and *M. psittaci* mitogenomes, which do not possess LIRs, G4s were located within the intergenic regions. Interestingly, only phylogenetic clades I and II incorporate G4DNA motifs in *trn*P, and their distribution is conserved within each of these clades. Potential G4s were also observed (a) in *cox*1 third intron (group-IB) of *M. sympodialis* and *U. maydis*, (b) in *atp*9 of the early diverging species *M. slooffiae* which is not part of its unique LIR, and (c) in the *trn*K of *M. caprae*.

**Figure 4.**
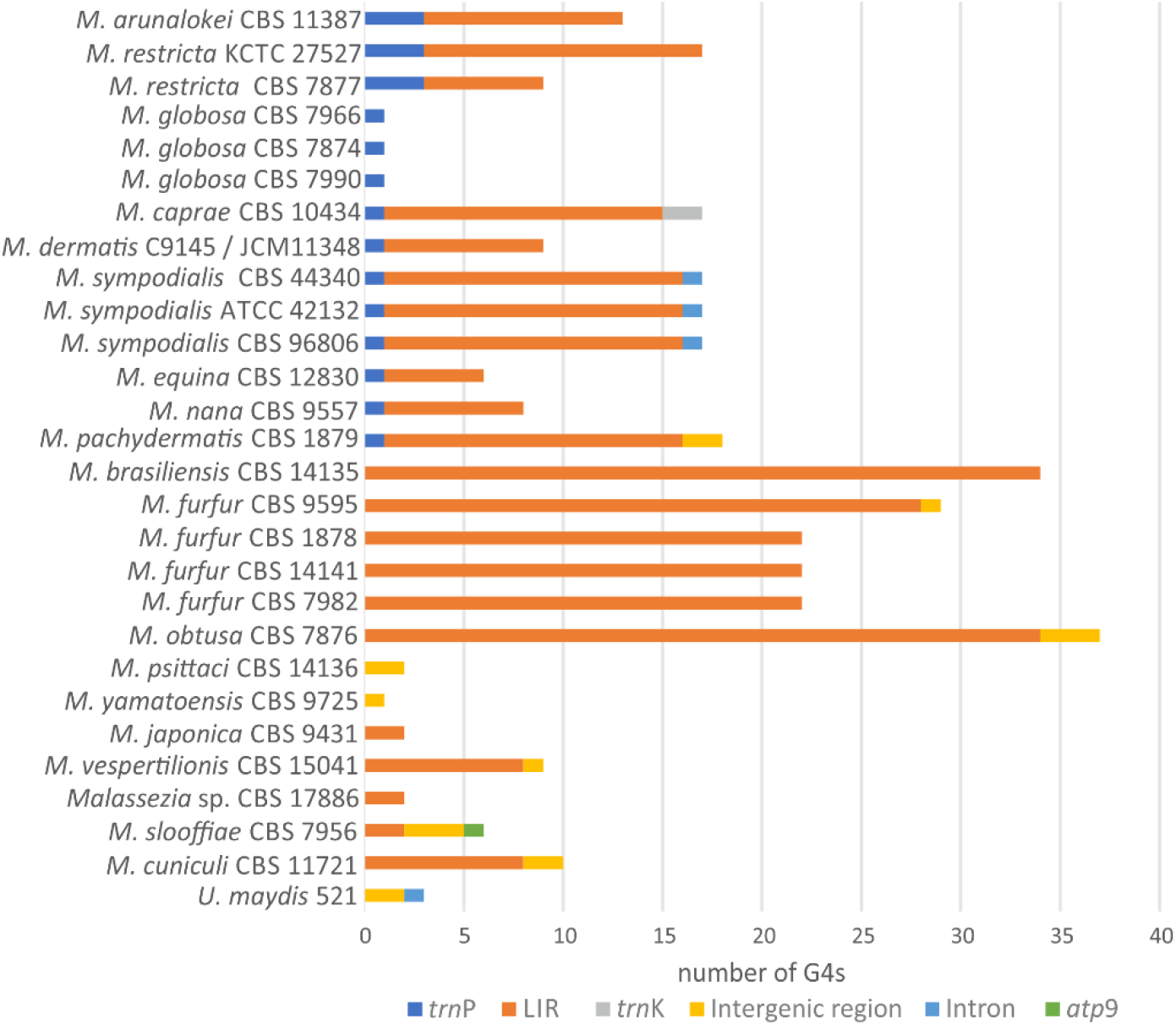
The G-quadruplex distribution in *Malassezia* mitogenomes. The G4s located in the *trn*P, LIRs, *trn*K, intergenic regions, introns and *atp*9 gene are shown with blue, orange, grey, yellow, light blue and green color, respectively.

## Discussion

The mitochondrion is a vital organelle for cell growth and survival of eukaryotic cells including fungi, but recent research has also implicated mitochondrial functions in the ability for pathogenic fungi to cause disease^29,30^. Mitochondrial genomes may also provide useful insights into fungal evolution as previously shown^4,5^. In general, whole genome sequencing projects focus on the nuclear genomes and thus far, only 192 out of 1,189 available fungal mt genomes belong to Basidiomycetes (https://www.ncbi.nlm.nih.gov/genome/browse#!/organelles/). However, species of Basidiomycetes present a complex mitogenomic gene organization with both ancestral and newly acquired properties, rendering them excellent model organisms for studying fungal evolution. Additionally, the medically relevant genus *Malassezia* embodies scarcely observed genetic elements, like LIRs, in their mitogenomes^3,28^. In this study, the mitogenomes of 11 *Malassezia* species along with publicly available mitogenome data (totaling 28 mt genomes, representing 18 described species and two putative new species) were utilized in a comparative genome analysis of all species of the genus *Malassezia*, in order to unravel the evolution of its mitogenome.

As expected, all *Malassezia* mt genomes contain 14 protein-coding genes that are present in nearly all fungi and metazoa, plus the *rps*3 gene which is present only in ca. 65% of Basidiomycetes^31^. *Malassezia* mitogenomes demonstrate substantial variation among species, mostly resulting from variation from intron size and frequency, intergenic regions, and LIR diversification, leading to mt genome sizes ranging from 28 kb to 64 kb.

A recent study presented a whole nuclear genome based phylogenetic tree for 15 out of the 18 currently described species^32^ and identified four main phylogenetic clusters (A-D) that correspond well with the mitochondrial topology of this work^33^. Here, the additional inclusion of *M. brasiliensis, M. psittaci, M. arunalokei*, completed the phylogenetic status of the whole genus from a mitochondrial perspective. The phylogenetic positioning of these three taxa is in agreement with recent publications concerning species barcoding based on nuclear data^34,35^, showcasing the value of concatenated mitochondrial PCG matrices in fungal phylogenetics as was recently also shown for the subphylum Saccharomycotina of Ascomycota^5^.

In fungi, variation in gene order is immense^36–38^. Even when only considering the genus *Malassezia*, a diversity in mt genome organization was observed (Figure 1). Only six shared syntenic blocks exist (Figure 2), of which *nad*2-*nad*3 seems to form a syntenic block in most fungal species^39^. The only other two genes that generally exist next to each other in fungi are *nad*4L-*nad*5 and it has been suggested that the variations in gene order is most likely the result of recombination^5,37^. Gene organization seems to be more discrepant between basal species and has a tendency to stabilize in the lately evolved phylogenetic clades.

All but two *Malassezia* species contain a LIR of variable size and content (Figure 3). Even the mitogenome of the earlier diverging species *M. cuniculi*, contains a unique inverted duplication (Figure 3). Another, evolutionarily independent species-specific duplication happens in *M. slooffiae*, which is the only species that additionally contains the *rnp*B gene which is uncommon in Basidiomycetes. This gene encodes the RNA subunit of mitochondrial RNaseP, an enzyme necessary for the maturation of the tRNAs^37^. It has been shown that *rnp*B is a functional gene in Ascomycetes and the protists *Reclinomonas americana* and *Nephroselmis olivacea*, and it is considered an ancestral element which was probably lost in most fungal species during evolution^40^. However, the existence of this gene in the intra-IR region of *M. slooffiae* may suggest a evolutionarily more recent addition to the mitogenome due to the LIR mobility. The LIRs of the remaining species seem to be the result of a continuous evolutionary process of three steps, starting with the duplication and inversion of the ancestral syntenic unit *trn*H-*trn*M in the mitogenome of *Malassezia* sp. CBS17886 (Figure 3, step 1). As a second step of the process, the duplication of ancestral syntenic unit *trn*R-trnL-*atp*9, also found in *U. maydis* (NCBI Acc No. NC_008368.1) and *Pseudozyma* sp. (NCBI Acc No. MK714018.1), and the loss of the one *trn*H copy led to the formation of the LIRs in phylogenetic clades I, II and in *M. vespertilionis*. Subsequently, in the other species of clade III an inversion of the previous syntenic unit is observed along with the deletion of one of the two *trn*Ms, resulting in an intra-IR region containing only *trn*M-*trn*H. In agreement with previous studies, the mechanism might have been recombination^41^ which in this case leads to a cross-over fixation, resulting in the loss of *trn*M. Interestingly, between closely related species with comparable LIR composition in clades I, II and III, the intra-IR is detected in normal or inverted orientation. Studies focusing on intra-species variation confirmed the presence of multiple mt genome rearrangements in various strains of one species, resulting from recombination between LIRs^3,26^. This could possibly apply for all *Malassezia* species but requires further exploration. The LIR variability in *Malassezia* is most probably the main course for the diverse synteny seen within the genus. This hypothesis is in agreement with the work of Liu and colleagues (2020) who demonstrated the presence of two different mtDNA genotypes in the same cell of *Agrocybe aegerita*, as a result of intramolecular recombination^42^. Up to date, LIRs have only been described for a few fungal genera, including several *Candida* species^28,41,43^, *Saccharomyces cerevisiae*^44^, *Hanseniaspora* sp. (*Kloeckera africana*)^45^ and several species of Agaricales^46^. Mito-LIRs can be found in a certain array of “lower” eukaryotes, like the oomycetes *Achlya ambisexualis*^47^ and the protozoan *Tetrahymena pyriformis* in which the mitogenome is linear^48^.

Previous studies showed that LIRs are recombination hotspots leading to the initiation of replication in *Candida albicans*^41^. A comparative study exploring the mt genomes of eight *Candida* species determined that LIRs may be involved in the so-called flip-flop recombination resulting in genome isomers and it was demonstrated that LIRs are involved in conversions between circular and linear mitochondrial topologies and in extent they may play a role in replication^43^. A possible mode of LIR introduction into mitochondria is plasmid integration^49^, although this study does not confirm this mechanism of origin for *Malassezia*, since no DNA polymerase gene or other plasmid related elements were found in all species examined. The lack of LIRs in two *Malassezia* species indicates that replication is not performed due to the LIRs. Finally, the LIR variation with its expansion and/or contraction within the genus *Malassezia* suggests that it is an ongoing process with back and forth results as observed in the different species.

In general, fungal mitogenome analyses present considerable genome size variation, which until now has been contributed to their sequence alterations resulting from mutation and recombination^5,50^. On the other hand, in plants and more specifically in chloroplast genomes, size diversity may be explained by sequence complexity as well as by the amount of repeated DNA^51^. Chloroplast genomes (cpDNA) usually present a commonly found structure of two single regions divided by LIRs^52^ allowing cpDNA to evolve under strong constraint -either mechanistic or selective – in order to maintain a compact, largely genic genome^51^. In fungi, the lack of LIRs in the majority of the species indicates a free evolution of the mitogenomes with respect to their size, with other proposed underlying mechanisms^37,53^.

Additionally, the existence of inverted repeats is strongly related to the high number of G4s in the *S. cerevisiae* mitogenome^8,11^. In the current study, a similar observation is made for the mitogenomes of the genus *Malassezia* (Figure 4). In species which do not possess LIRs, G4 formation can occur in intergenic regions, *trn* genes and in some cases within introns (Figure 4). Interestingly, G4 structures were overlapping with the *trn*P gene in *Malassezia*’s later diverging phylogenetic clades I, and II. This phenomenon may be additional evidence to the hypothesis of *trn* genes acting as recombinational hotspots for mitogenome gene shuffling^5,54^. Moreover, this distribution contradicts with the respective G4 analysis in *S. cerevisiae*, where G4s did not overlap with *trn*s^11^. G4 DNA structures have already been related to DSBs^9,55^. Their location within the LIRs, which have been implicated in recombination events whenever DSBs exist, offer a repair mechanism and contribute to genome stability, despite the G4 DNA implication to deleterious effects on mtDNA^55^.

The following hypothesis concerning fungal mitogenome evolution may be suggested: In contrast to the cpDNAs, mitogenomes rarely contain cpDNA-like structures, and whenever found in fungi they are tandemly scattered, i.e., in different orders/subphyla which are also phylogenetically distant. Thus, their acquisition must be a recent event which happened several times during fungal evolution, as phylogenetically scattered species carrying LIRs in their mt genome suggest. The G4 DNA structures seem to have coevolved with the LIRs, whenever both exist. In other words, they coexisted to act as a unit performing recombination and repair to stabilize the mitogenome.

In conclusion, *Malassezia*’s mitogenome analyses showed the existence of newly acquired elements like LIRs and G4 DNAs, which converged and provided a reliable recombination-based mechanism to facilitate genome stability. This work adds to our understanding of the mechanisms that drive mitogenomic evolution.

## Materials and Methods

### *Malassezia* species and strains

In this study 28 *Malassezia* strains, corresponding to all known species of the genus, including a putative new species from bat, were used in the *in-silico* comparative mt genome analysis. In detail, 11 mt genomes were retrieved from in house generated Whole Genome Sequencing data (WGS) (Table 1).

### Culture and DNA extraction

*Malassezia* strains were grown on modified Dixon agar^56^ for 48 – 96 h at 30°C (*M. cuniculi* on Leeming and Notman agar for 7 d at 36°C; CBS 17886 for 7 d at 24°C), and cells were harvested into 50-mL tubes. Yeast cells were lysed, following the Qiagen genomic DNA purification procedure for yeast samples (Qiagen, Hilden, Germany), with some modifications. Lyticase incubation was performed for 2 h at 30°C, and RNase/Proteinase incubation was performed for 2 h at 55°C. Genomic DNA was purified using Genomic-tip 100/G prep columns, according to the manufacturer’s instructions.

### Whole genome sequencing

Samples were sequenced on the Oxford nanopore PromethIon platform, and libraries were prepared using the standard Native Barcoding Genomic DNA Ligation Protocol. Base calling was performed on the PromethIon (Release 21.02.7) with high accuracy setting.

### Long read assembly

De novo assembly was carried out by Flye version 2.9^57^. The “--nano-raw” flag was used; genome size was set to 8 Mb and polishing iterations to 2.

### Mitochondrial DNA assembly

For most strains, mitochondrial DNA was found in multiple fragments in the assembled genomes. To retrieve it in one piece, the following process was followed: 1) raw reads were mapped against the assembled genome, using minimap2 version 2.17-r941, 2) reads that mapped against chromosomal fragments were discarded; 3) remaining reads were reassembled using both Flye 2.9^57^, setting “--asm-coverage” to 50, and canu 2.2^58^. Since reads were not 100% mitochondrial and the exact size of the remaining chromosomal fragments was not known, multiple assemblies took place for every strain. Genome size was set to 50 kb, 500 kb or 1000 kb, until the mtDNA was retrieved in one scaffold. Where Illumina data were available, mtDNAs were assembled using GetOrganelle v1.7.5 using recommended parameters^59^. The obtained mt scaffolds were additional validated by aligning them against known *Malassezia* mt genomes using tBLASTn/BLASTx/BLASTn^60^ and were further analyzed as described below.

### Mitochondrial genome annotation and comparative analyses

The mt scaffolds were annotated as follows: the protein coding and the ribosomal (rRNA) genes were identified using BLASTx and BLASTn, respectively. The tRNA genes were detected using the web-based tRNAScan-SE platform^61^. Intron characterization was performed by RNAweasel online tool^62^ and the intron borders were also verified manually. The open reading frames (ORFs) were identified using ORF finder (https://www.ncbi.nlm.nih.gov/orffinder/) setting “ORF start codon to use” parameter as “ATG and alternative initiation codons", and the genetic code employed was “The Mold, Protozoan, and Coelenterate Mitochondrial Code and the Mycoplasma/Spiroplasma Code” (NCBI transl_table=4). The 11 newly annotated mt genomes of this study were deposited to GenBank (acc Nos: ON585844-ON585854). The sequence of the nine conserved mt gene blocks identified in synteny analysis, were collected, and aligned by multiple sequence alignment program MAFFT using the E-INS-i method^63,64^ in order to identify the mt sequence diversity in *Malassezia* spp.

### Phylogenetic analyses

Amino acid sequences as produced by the 15 mt protein coding genes (PCGs) (i.e., *atp*6, *atp*8-9, *cob, nad*1-6, *nad*4L, *cox*1-3 and *rps*3) were collected from the 28 *Malassezia* mt genomes for phylogenetic purposes (Suppl. File S2). *Ustilago maydis* strain 521 was used as an outgroup (GenBank Acc. No. NC_008368) in all analyses besides the phylogenetic one. The sequences for each protein were aligned by Lasergene’s MegAlign v.11 program (10.1385/1-59259-192-2:71) using the ClustalW method with default settings. The concatenated dataset was created for phylogenetic purposes. The phylogenetic tree construction was performed using Neighbor Joining (NJ) and Bayesian Inference (BI) methods through PAUP4^65^ and MrBayes^66^ software, respectively. For NJ analyses, reliability of nodes was evaluated using 10.000 bootstrap iterations for all concatenated and individual datasets. For BI analyses, the determination of the evolutionary model, which was best suitable for each dataset, was performed using the program ProtTest (ver. 3)^67^. The BIC Information Criterion was applied, and the best nucleotide substitution model was found to be LG+I+G+F. Four independent MCMCMC analyses were performed, using 5 million generations and sampling set adjustment for every 100,000 generations. The remaining parameters were set to default.

### LIRs and G-quadruplex prediction

The LIR of each genome was identified in sequence level using Blastn against itself and the identical inverted repeats were collected. For the prediction of G-quadruplexes in *Malassezia*’s mitogenomes, G4Hunter web application were used setting the window in 25 bp and the threshold/G4Hunter score to 1,2^68^.

## Data availability

Mitogenomes have been submitted to the NCBI GenBank (https://www.ncbi.nlm.nih.gov/genbank/) under accession numbers: ON585844-ON585854.

## Acknowledgements

The authors wish to thank Prof. Dr. Ferry Hagen for his help with sequencing some of the strains and Dr. Jeffrey M. Lorch for providing isolate CBS 17886. Authors ACC and VNK would like to thank the Federation of European Microbiological Societies (FEMS) which nominated ACC the ‘research and training grant—FEMS-GO-2020–199’ for this project. This work was supported by “Special Account for Research Grants” of National and Kapodistrian University of Athens under Research Program (code no. 16673).

## Author notes

These authors contributed equally: Christinaki A.C. and Theelen B.

## Figures, tables and supplementary information

Supplementary Table S1. Detailed mitogenome characteristics in examined *Malassezia* species.

Supplementary Table S2. *Malassezia*’s mitochondrial introns.

Supplementary Table S3. Distance/similarity percentage matrix for the concatenated 15 PCGs.

Supplementary Table S4. G-quadruplex detailed analysis

Supplementary File S1. Schematic presentation of G-quadruplexes and their location in *Malassezia* mitogenomes.

Supplementary File S2. Concatenated amino acid matrix of the 15 mitochondrial PCGs in nexus format.

**Supplementary Figure S1.**
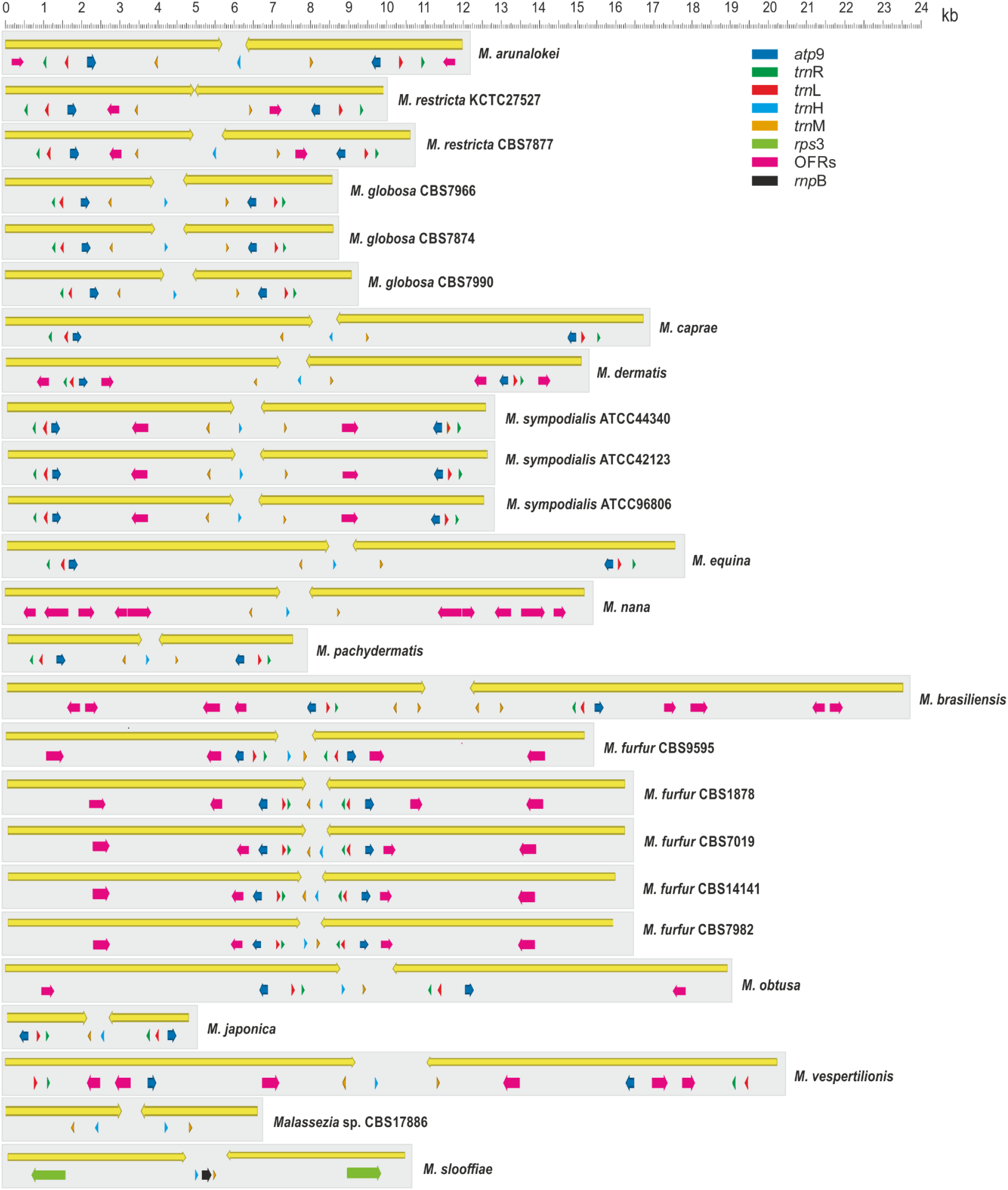
Detailed presentation of *Malassezia*’s LIR.

