## Supplementary material for "Co-evolution of Large inverted repeats and G-quadruplex DNA in fungal mitochondria may facilitate mitogenome stability: the case of *Malassezia*": Suppl. File S1

### *Malassezia arunalokei*

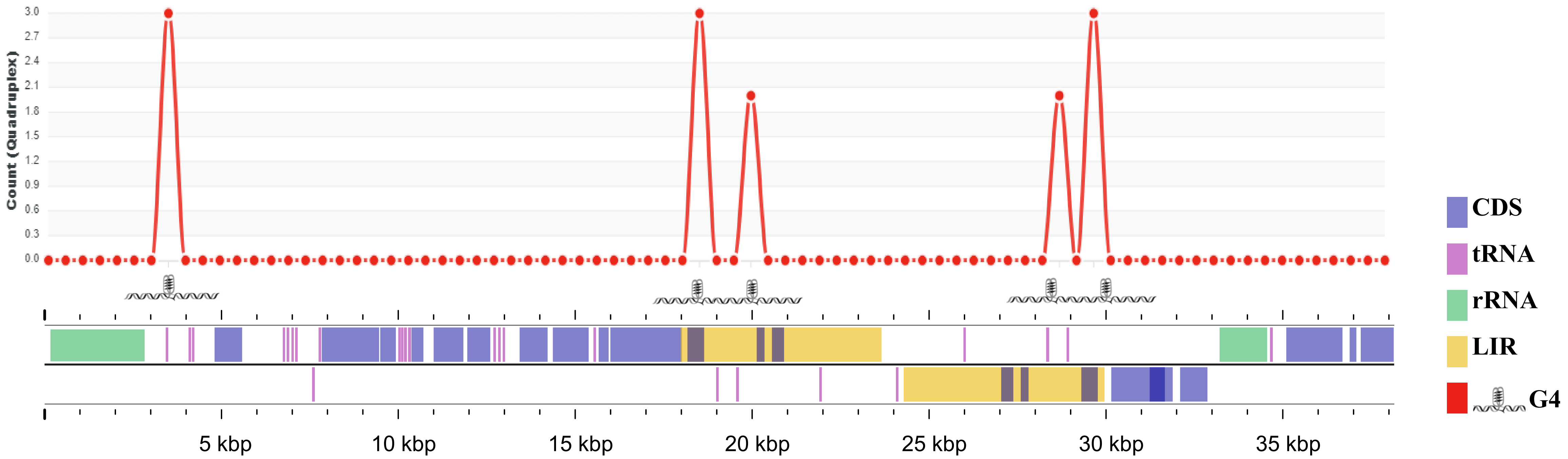

### *Malassezia brasiliensis*

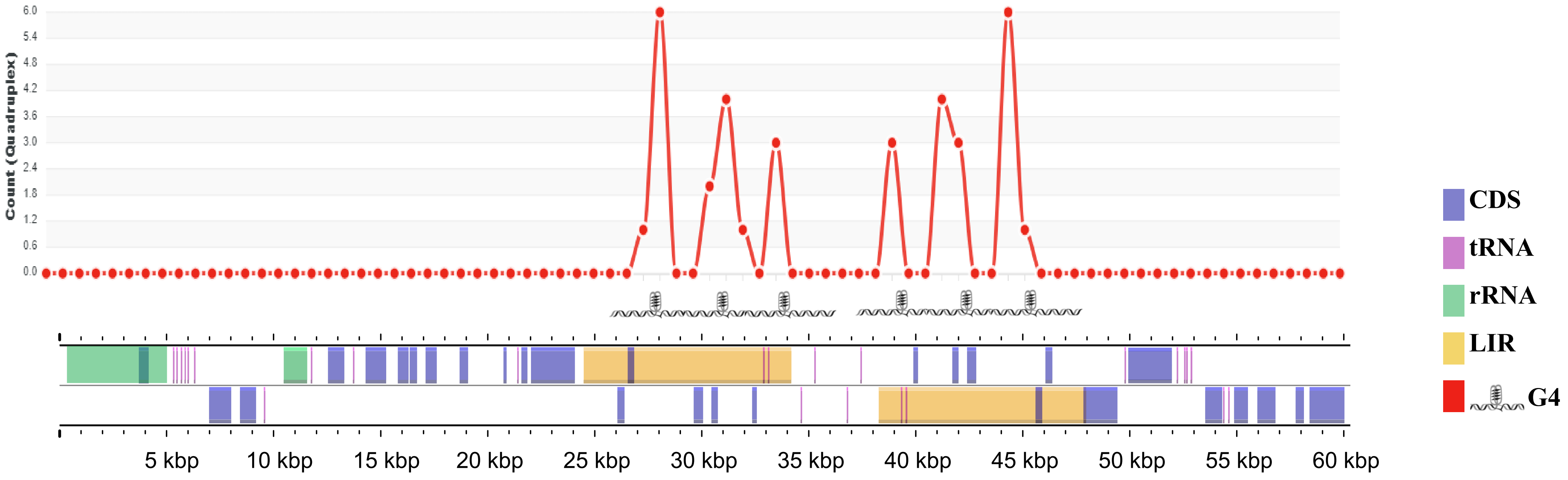

### *Malassezia caprae*

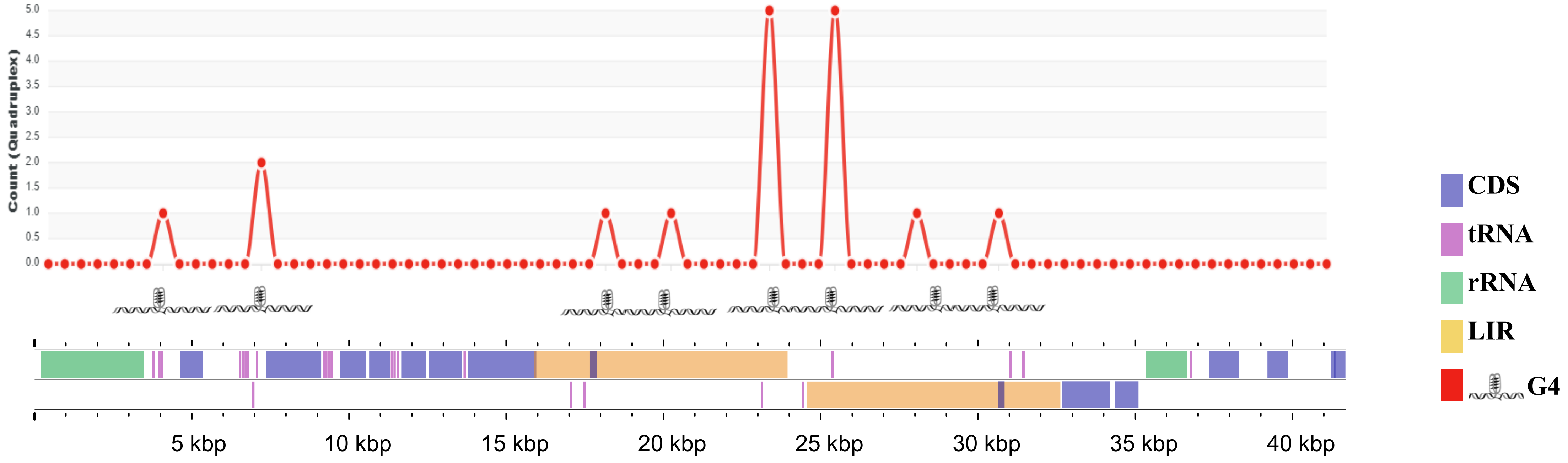

### *Malassezia dermatis*

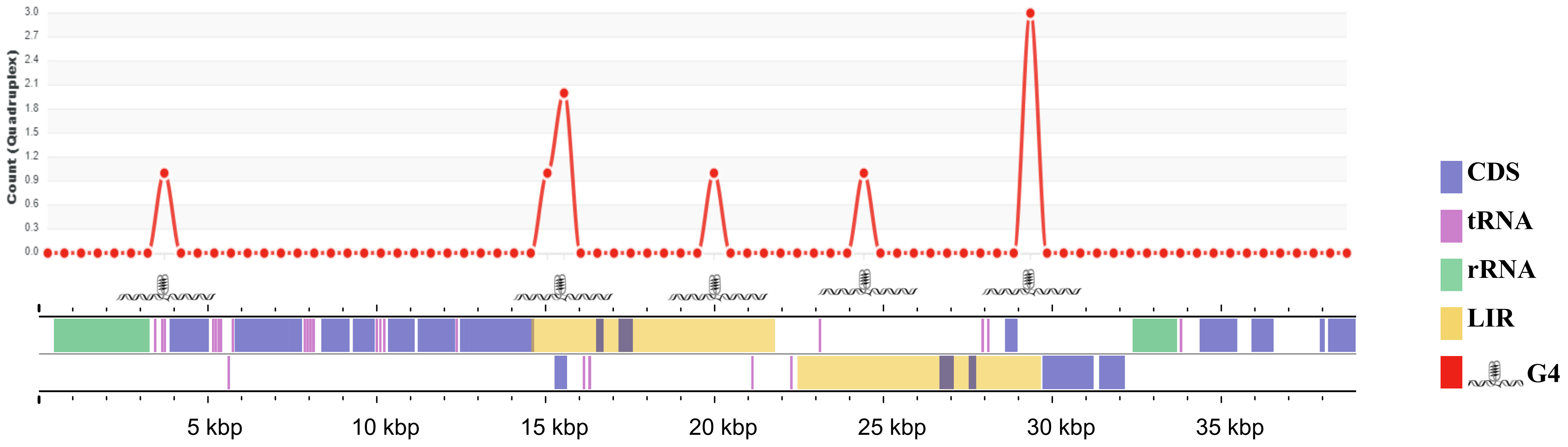

### *Malassezia equina*

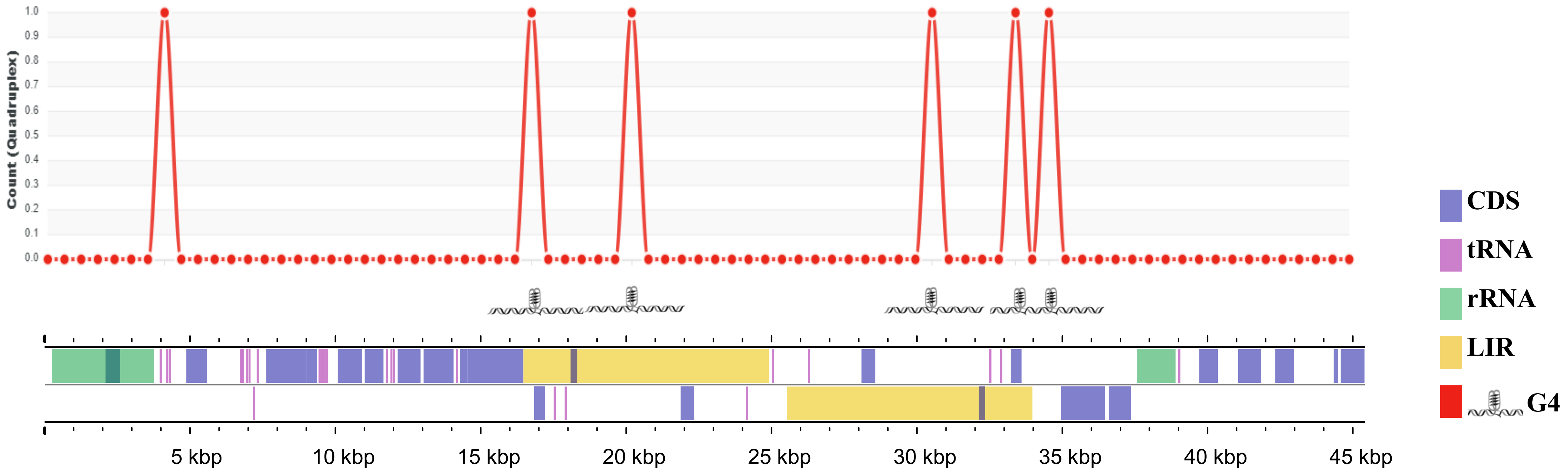

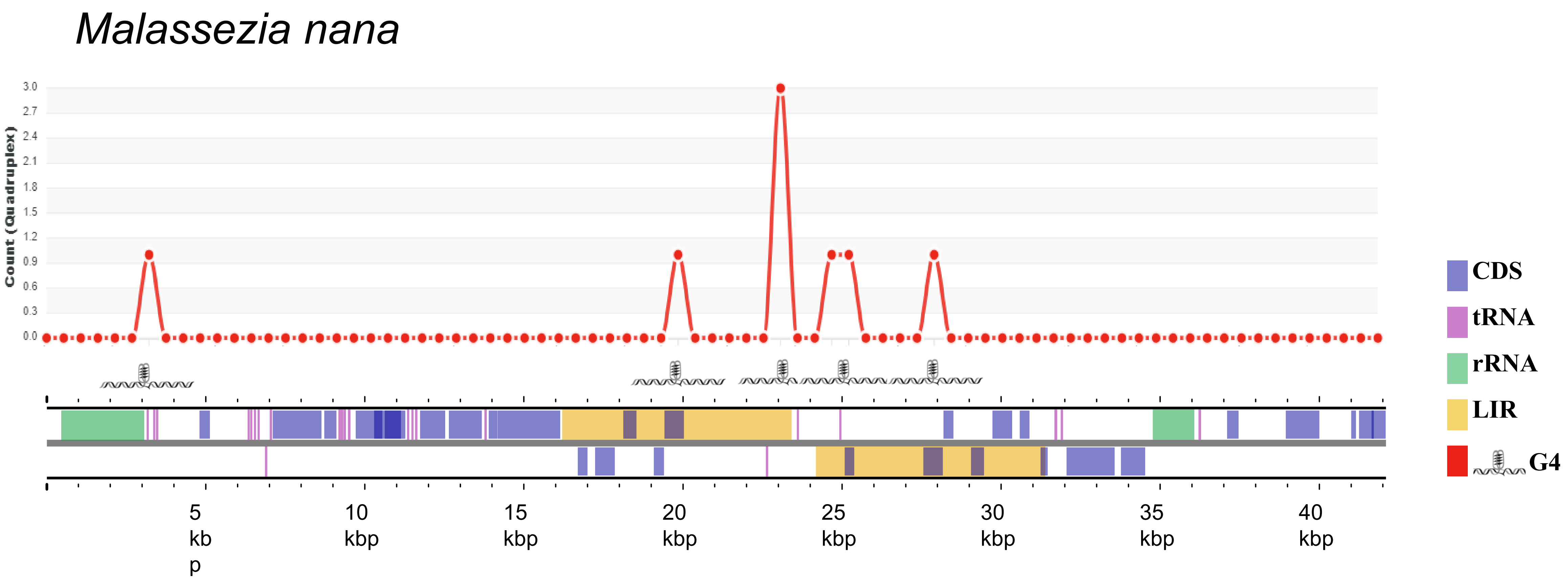

Malassezia nana
Count (Quadruplex)
3.0
2.7
2.4
2.1
1.8
1.5
1.2
0.9
0.6
0.3
0.0
CDS
tRNA
rRNA
LIR
G4
5 kbp
10 kbp
15 kbp
20 kbp
25 kbp
30 kbp
35 kbp
40 kbp

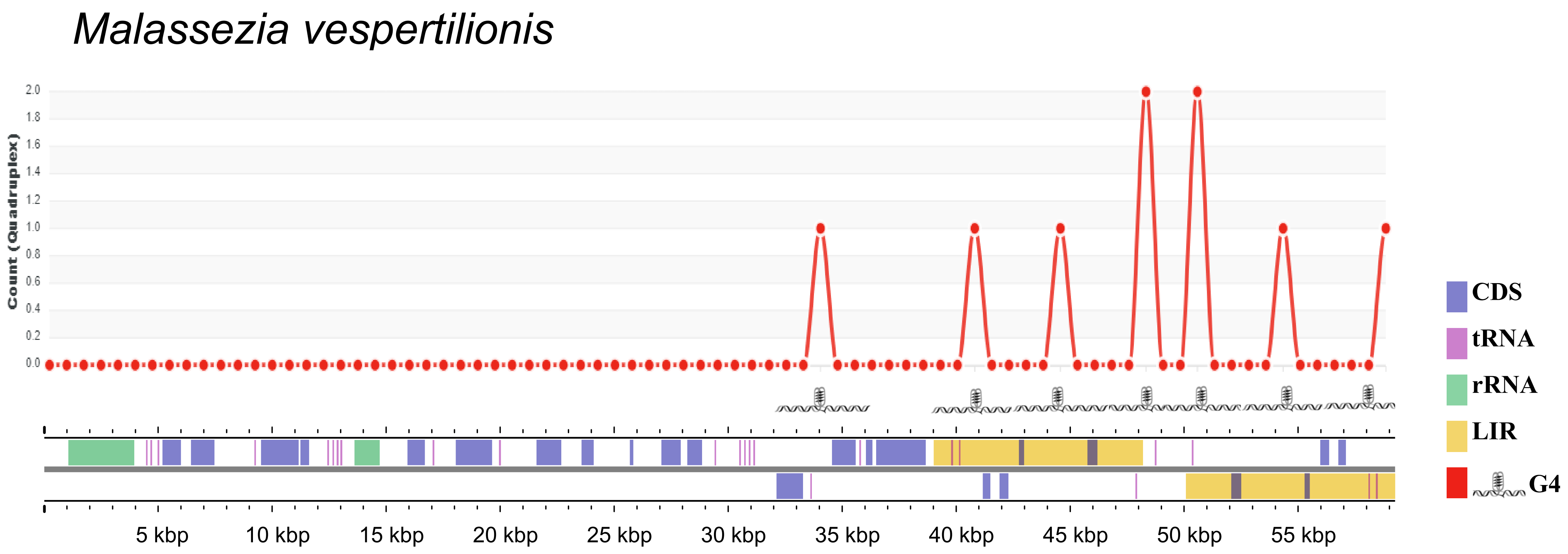

Malassezia vespertilionis
Count (Quadruplex)
2.0
1.8
1.6
1.4
1.2
1.0
0.8
0.6
0.4
0.2
0.0
5 kbp
10 kbp
15 kbp
20 kbp
25 kbp
30 kbp
35 kbp
40 kbp
45 kbp
50 kbp
55 kbp
CDS
tRNA
rRNA
LIR
G4

### *Malassezia spittaci*

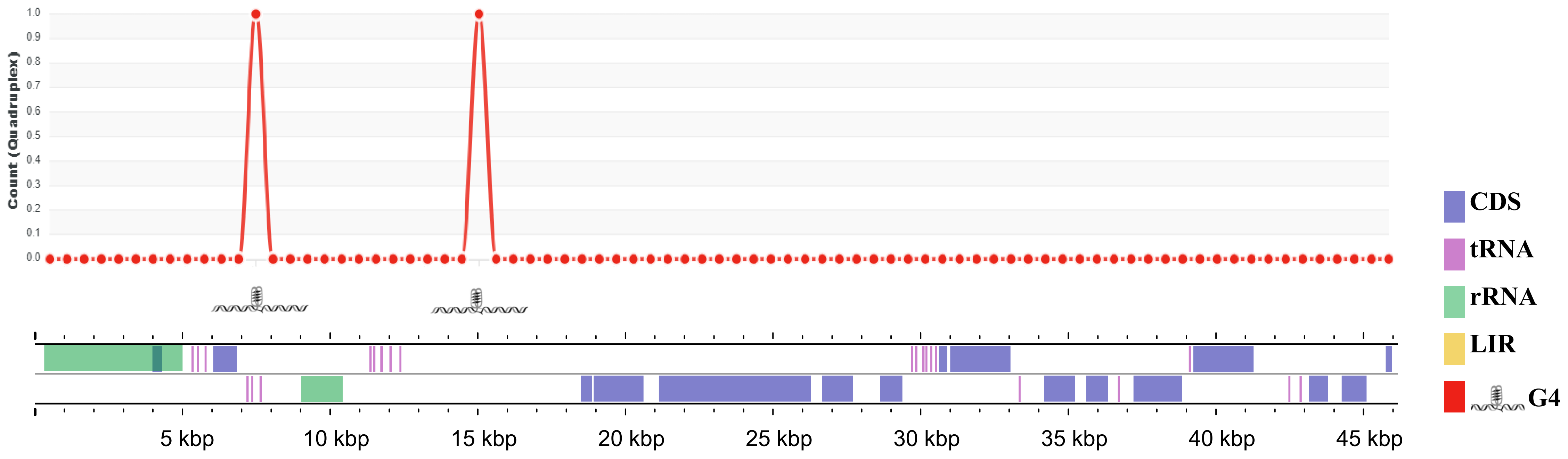

### *Malassezia* sp. strain CBS17886

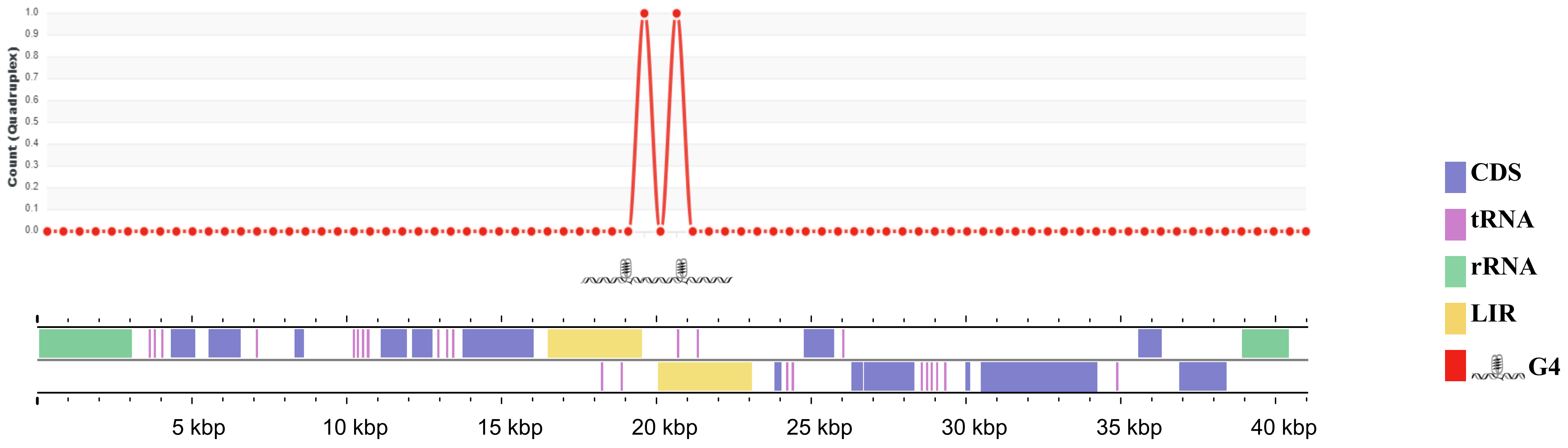

### *Malassezia furfur* strain CBS9595

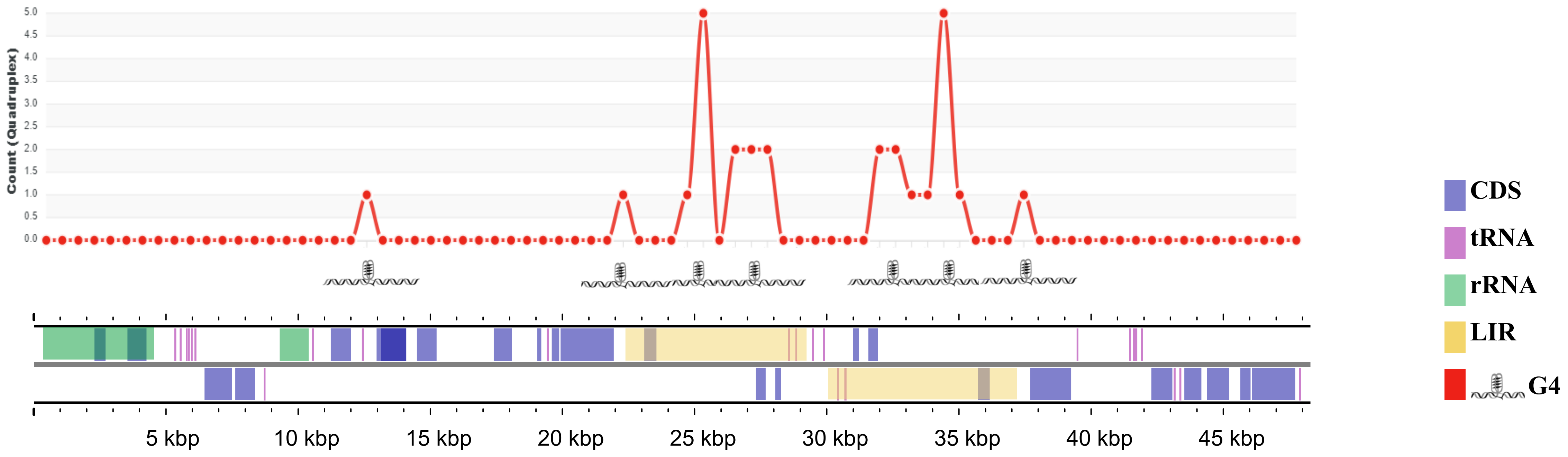

### *Malassezia yamatoensis* strain CBS9725

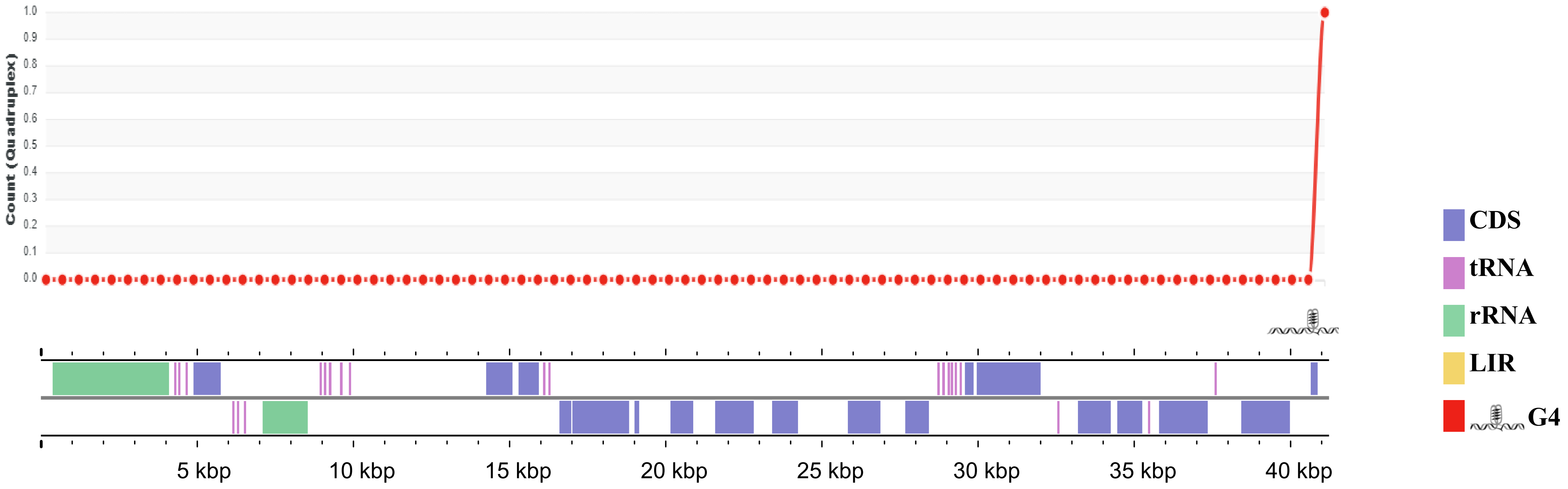

### *Malassezia furfur* strain CBS 1878

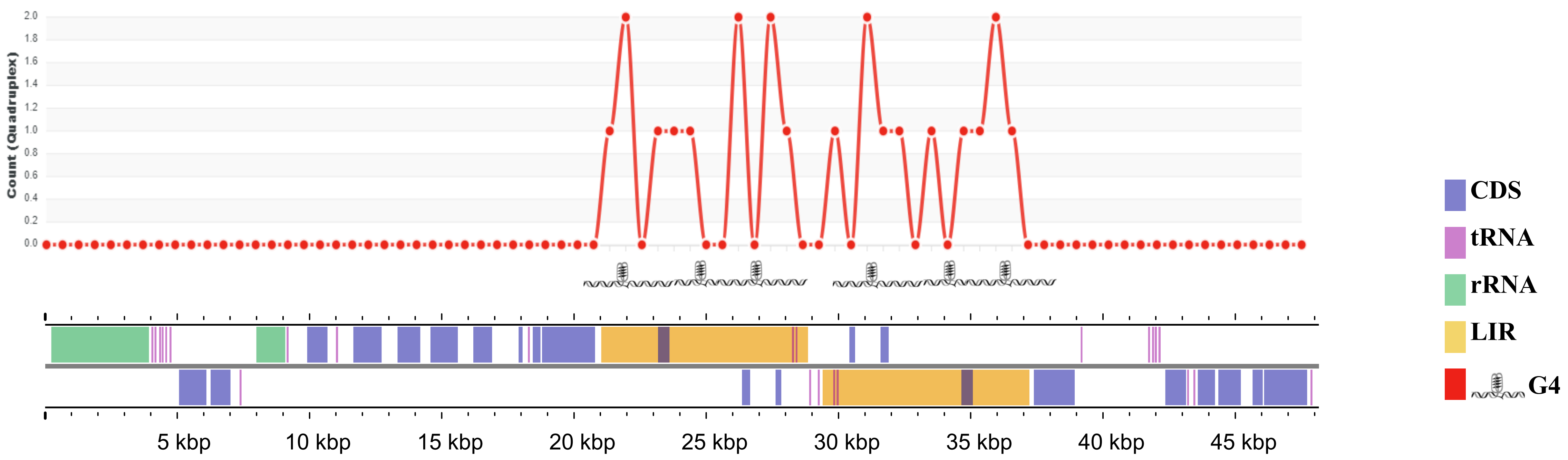

#### *Malassezia furfur* strain CBS 7019

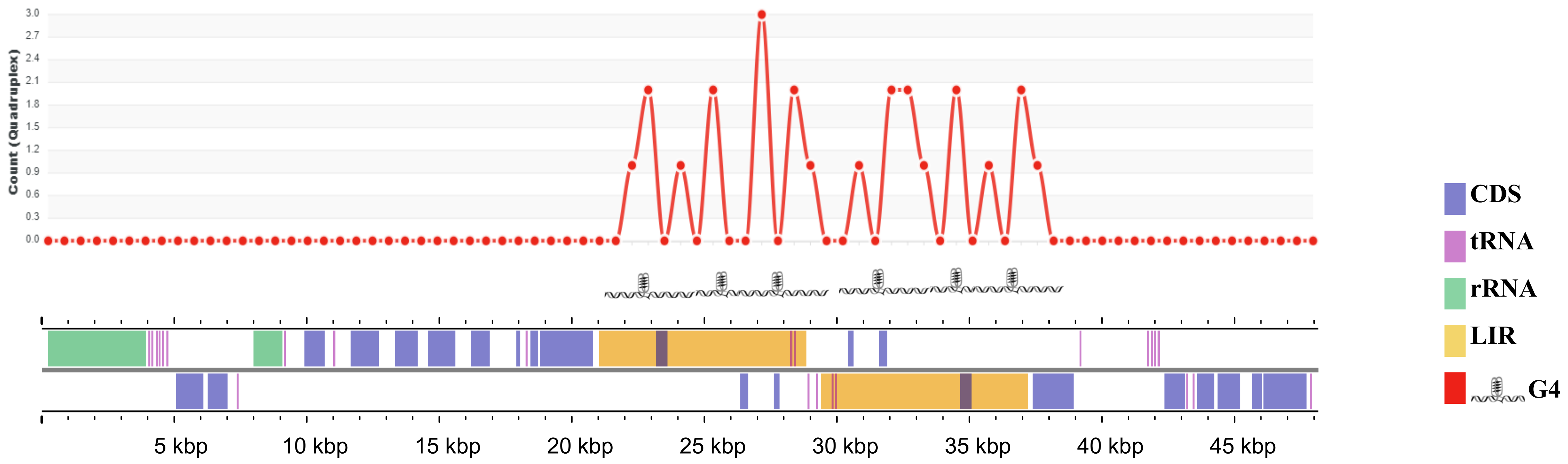

### *Malassezia furfur* strain CBS14141

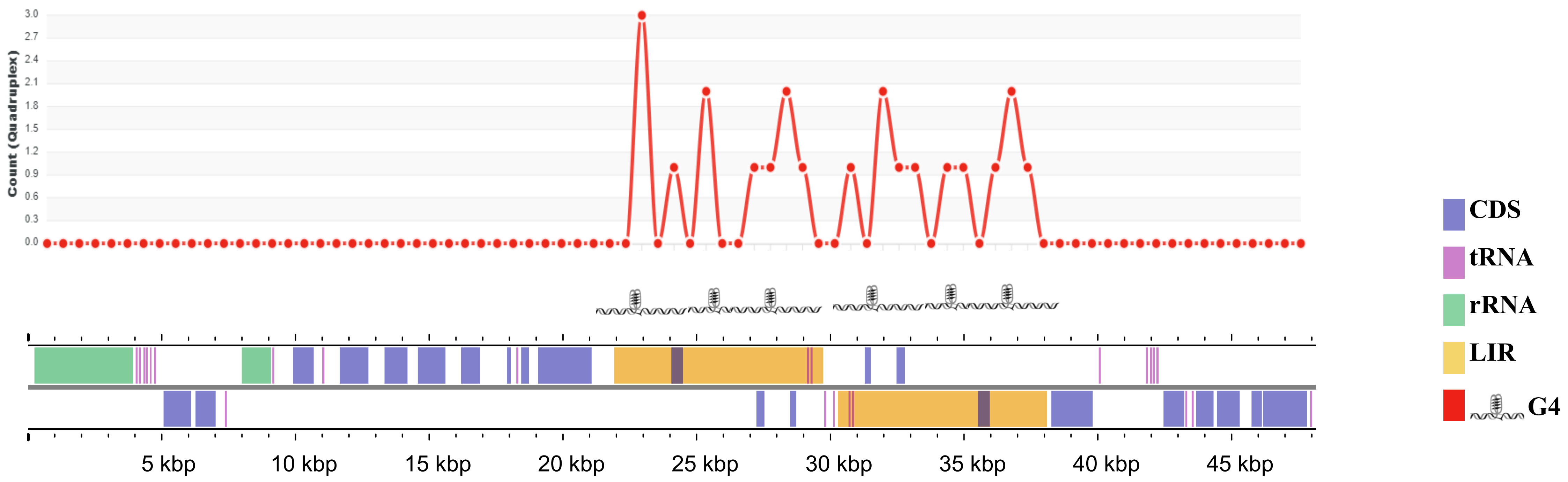

### *Malassezia furfur* strain CBS7982

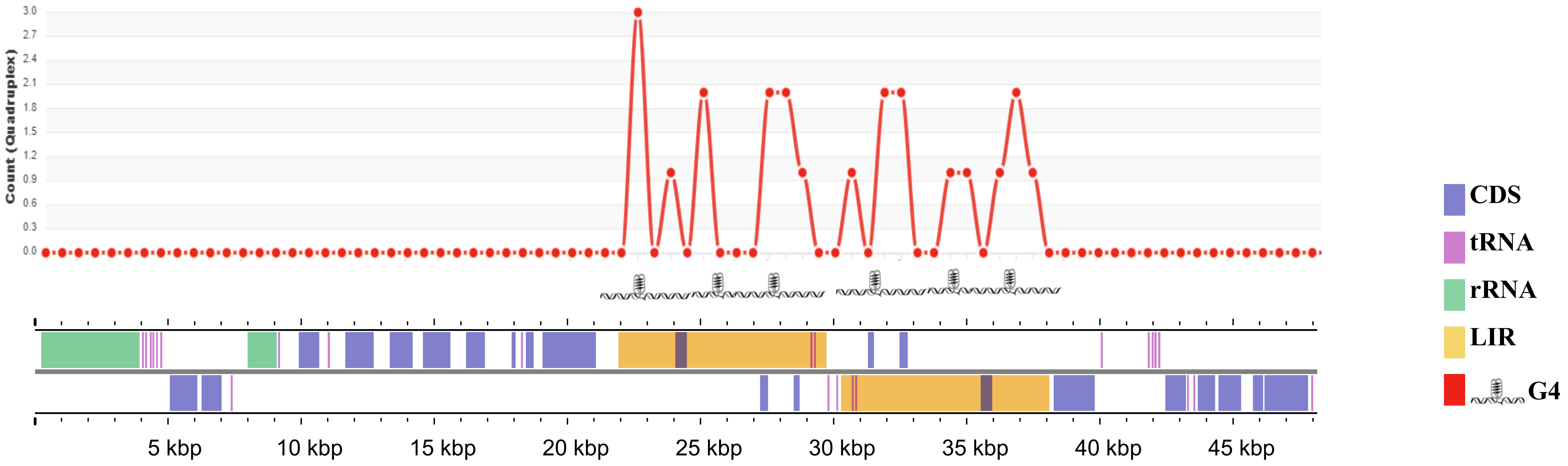

### *Malassezia obtusa* strain CBS7876

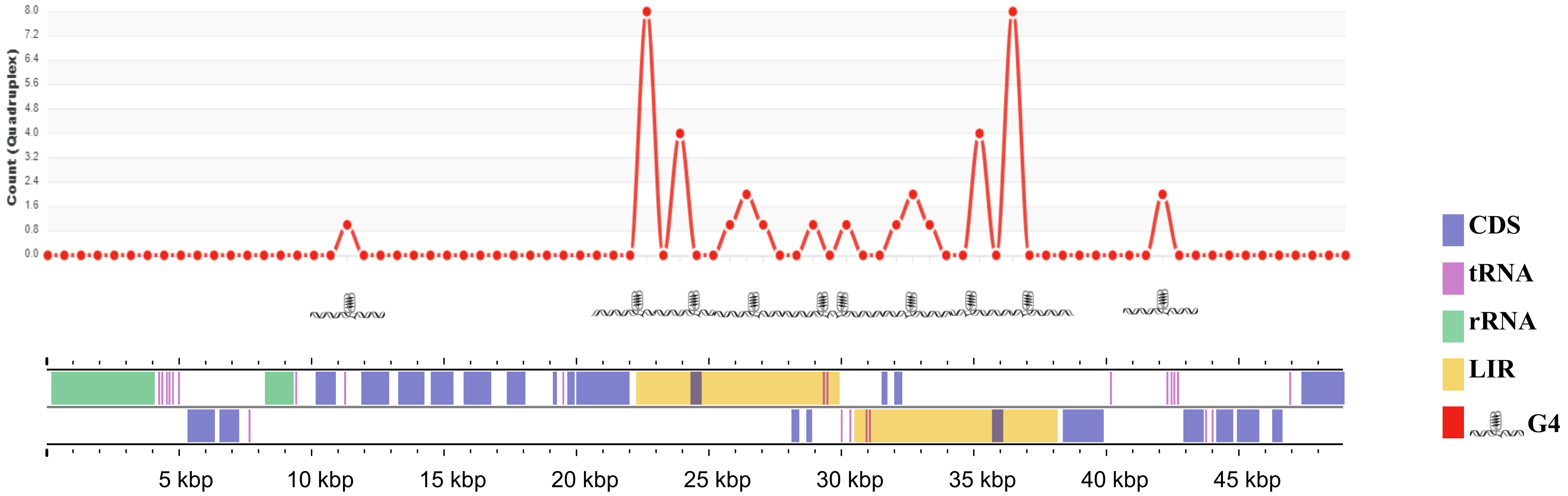

### *Malassezia japonica* strain CBS9431

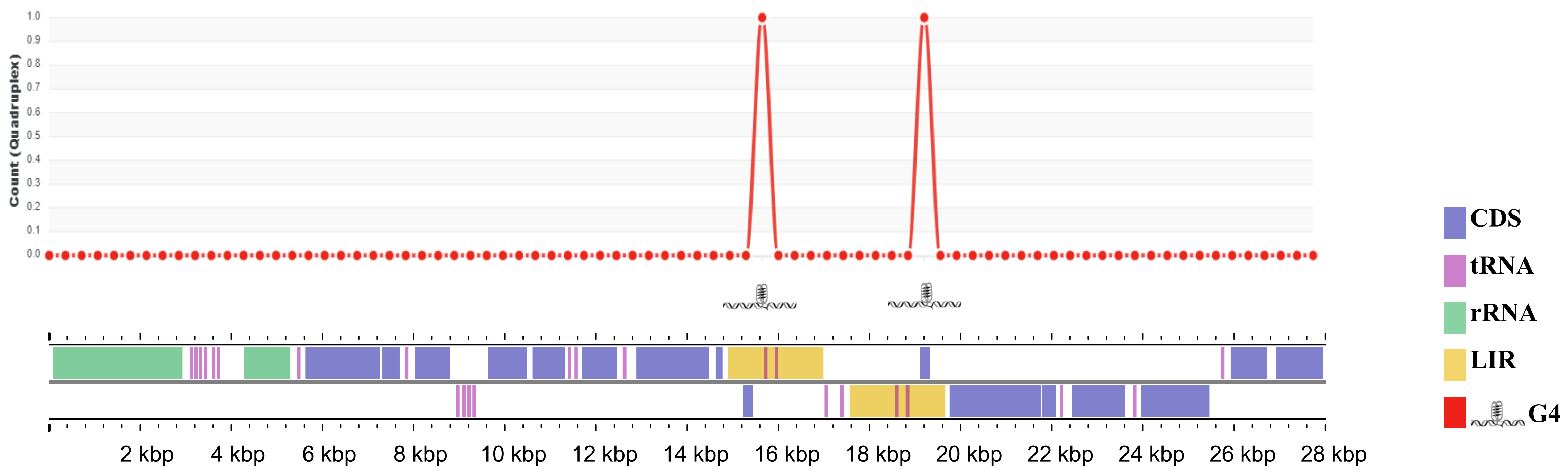

### *Malassezia restricta* strain KCTC 27527

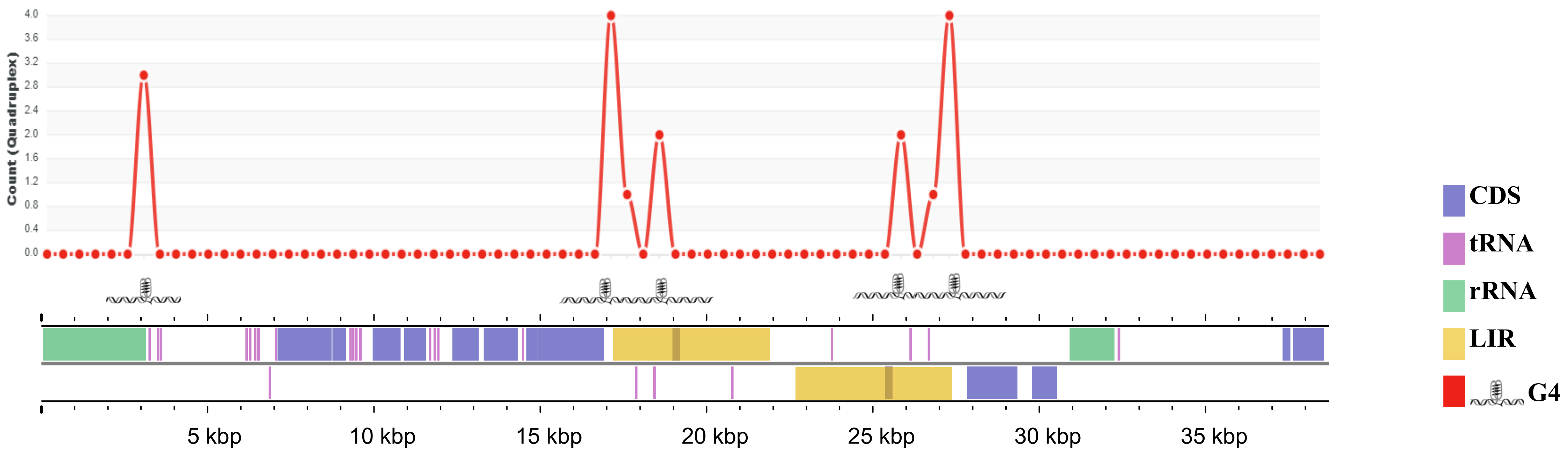

### *Malassezia restricta* strain CBS7877

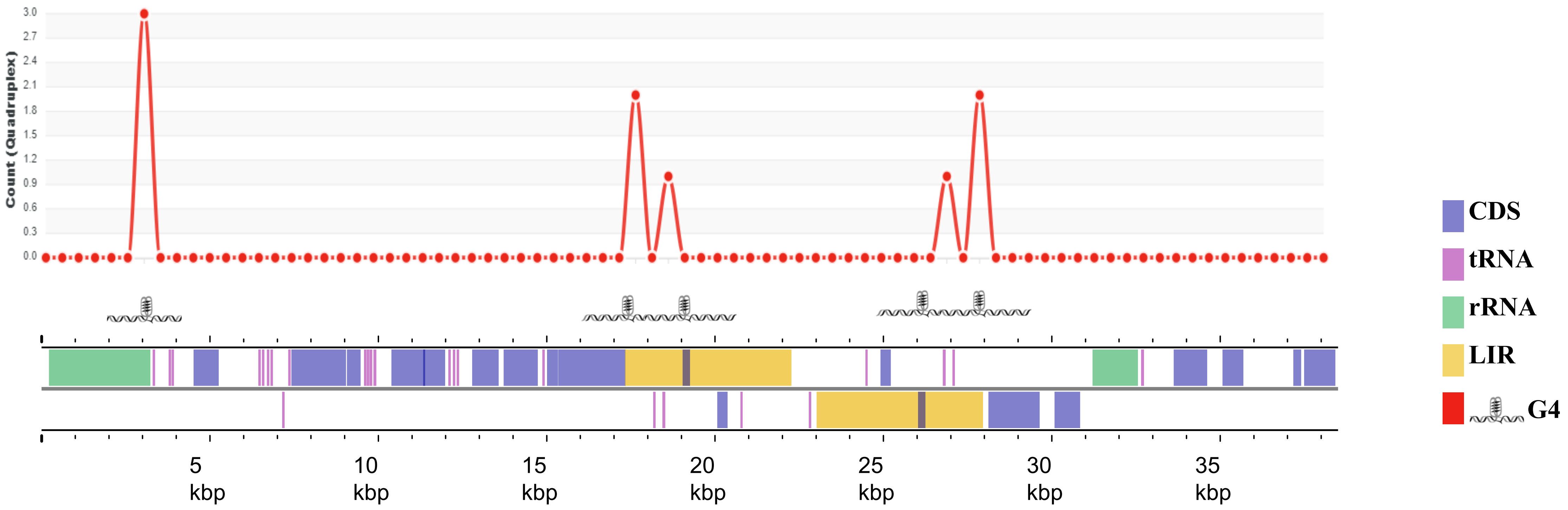

### *Malassezia globosa* strain CBS7966

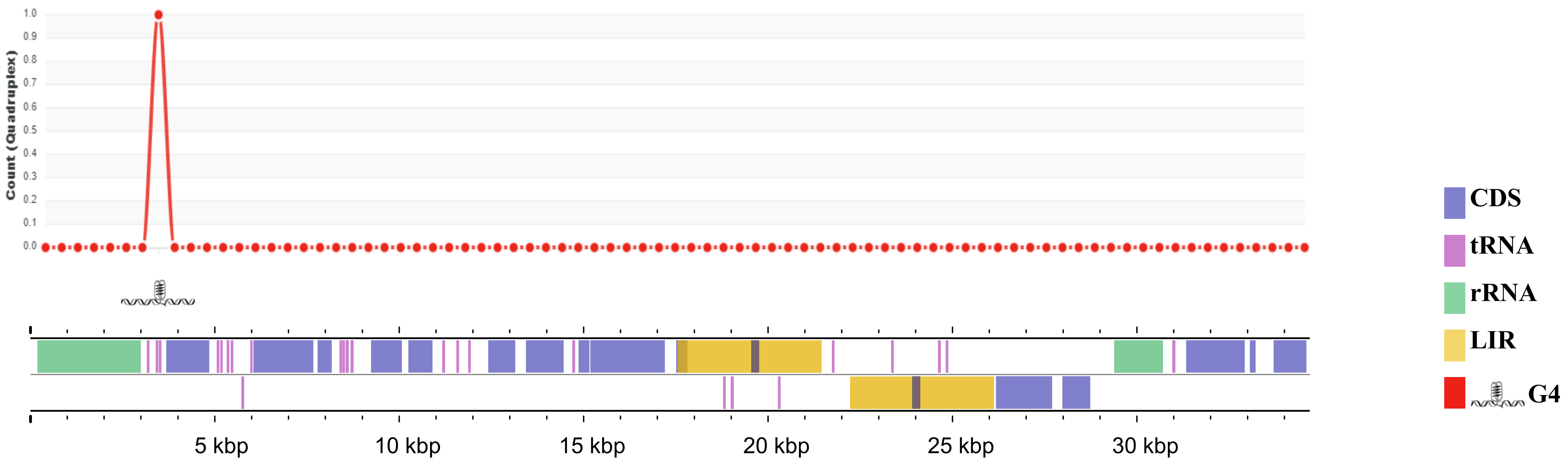

### *Malassezia globosa* strain CBS7990

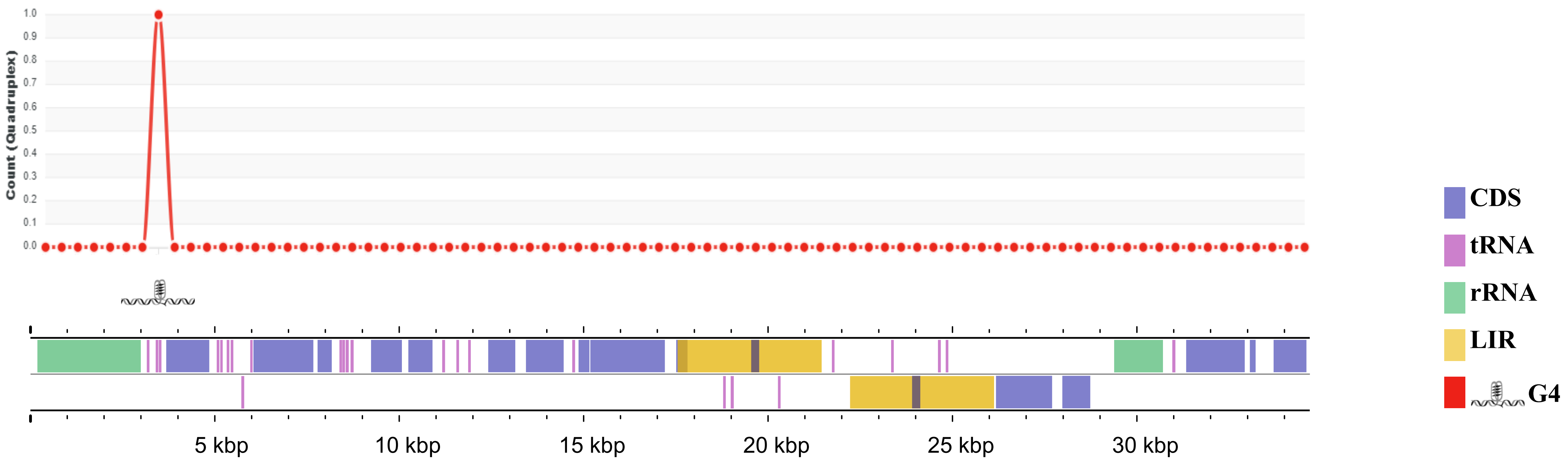

### *Malassezia globosa* strain CBS7874

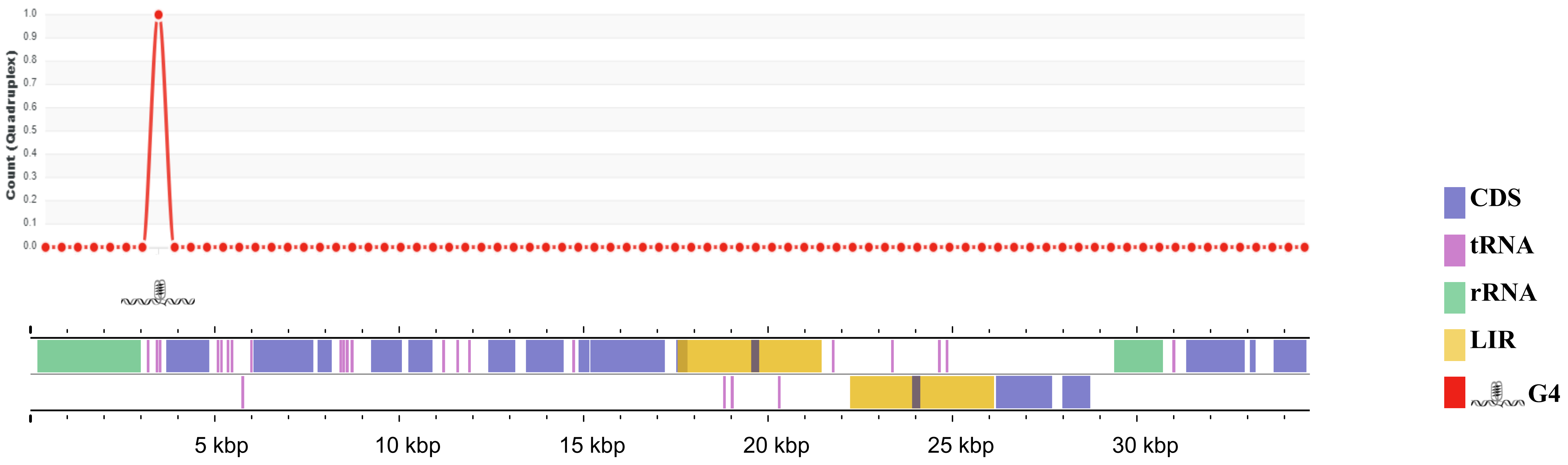

### *Malassezia sympodialis* strain CBS44340

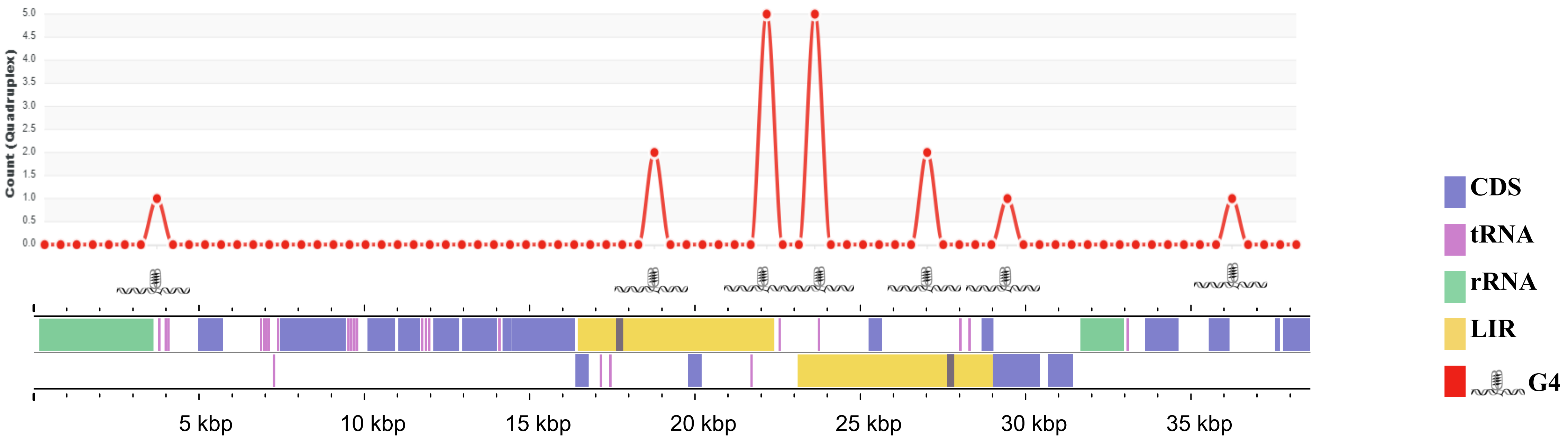

*Malassezia sympodialis* strain ATCC 42132

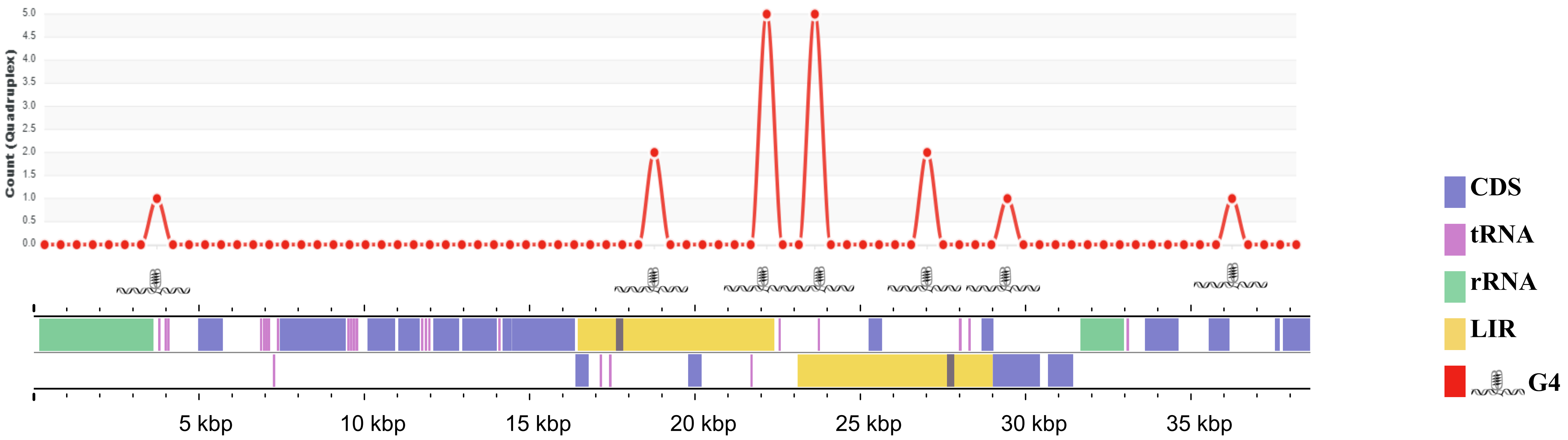

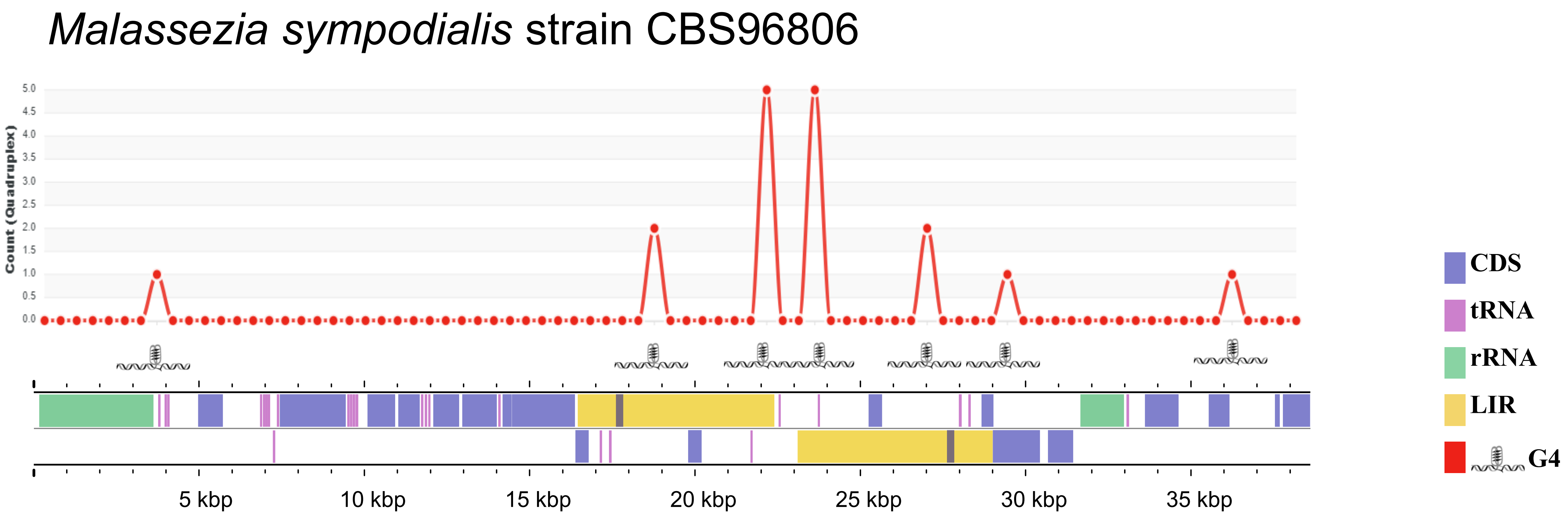

Malassezia sympodialis strain CBS96806
Count (Quadruplex)
5.0
4.5
4.0
3.5
3.0
2.5
2.0
1.5
1.0
0.5
0.0
5 kbp
10 kbp
15 kbp
20 kbp
25 kbp
30 kbp
35 kbp
CDS
tRNA
rRNA
LIR
G4

### *Malassezia pachydermatis* strain CBS1879

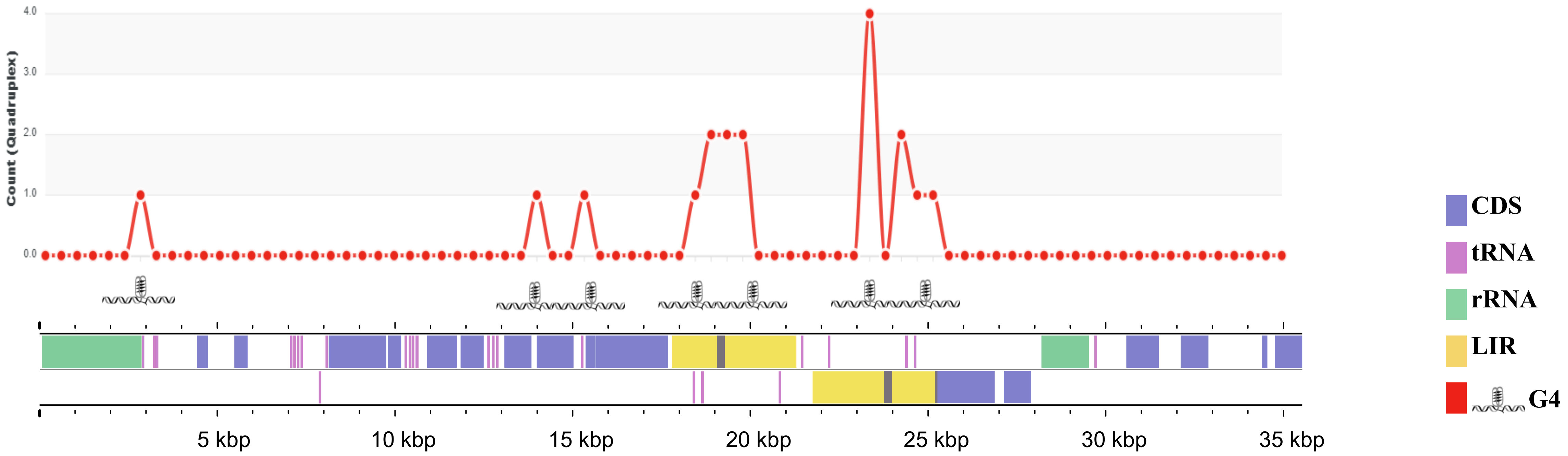

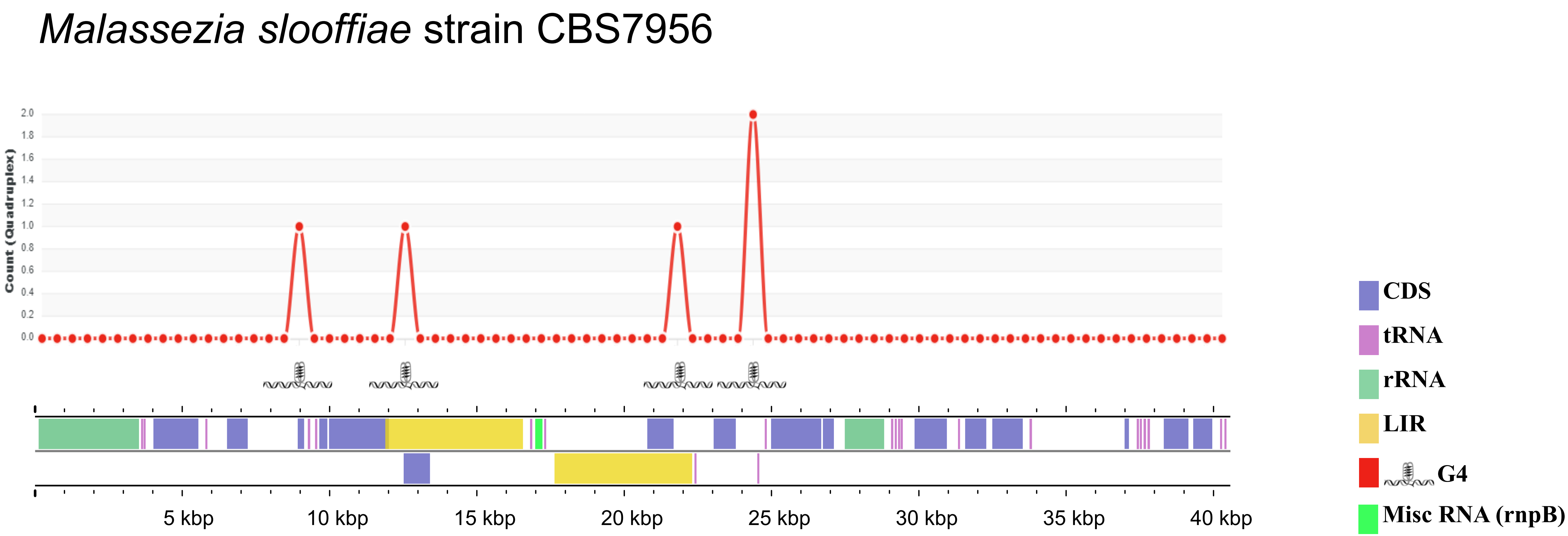

*Malassezia slooffiae* strain CBS7956
Count (Quadruplex)
2.0
1.8
1.6
1.4
1.2
1.0
0.8
0.6
0.4
0.2
0.0
5 kbp
10 kbp
15 kbp
20 kbp
25 kbp
30 kbp
35 kbp
40 kbp
CDS
tRNA
rRNA
LIR
G4
Misc RNA (rnpB)

### *Malassezia cuniculi* strain CBS11721

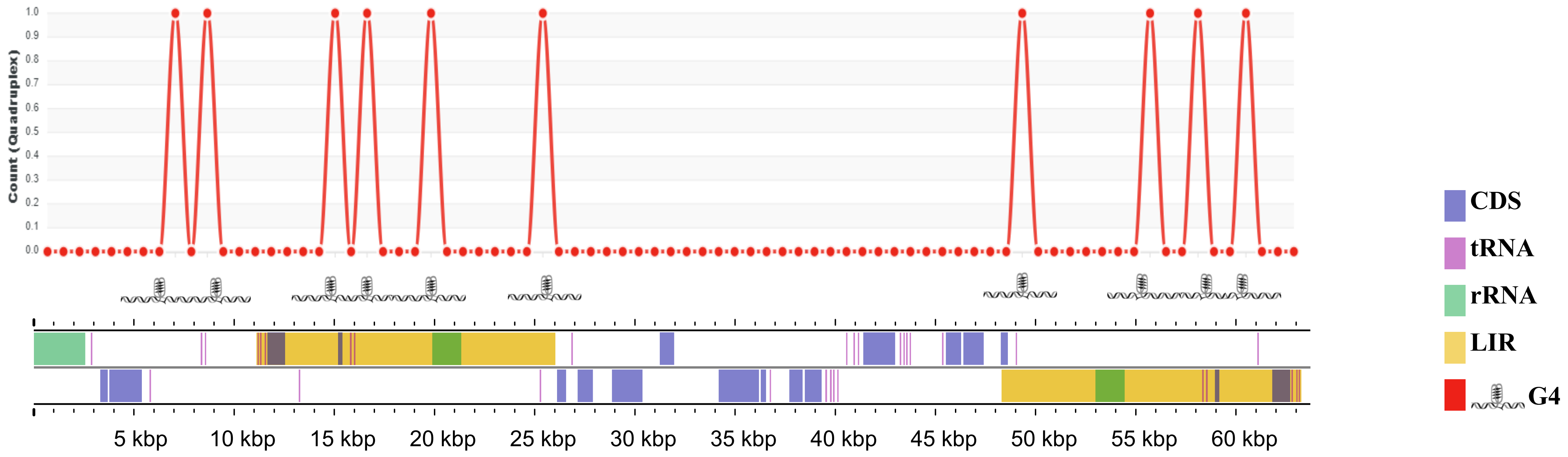

### Ustilago maydis strain 520

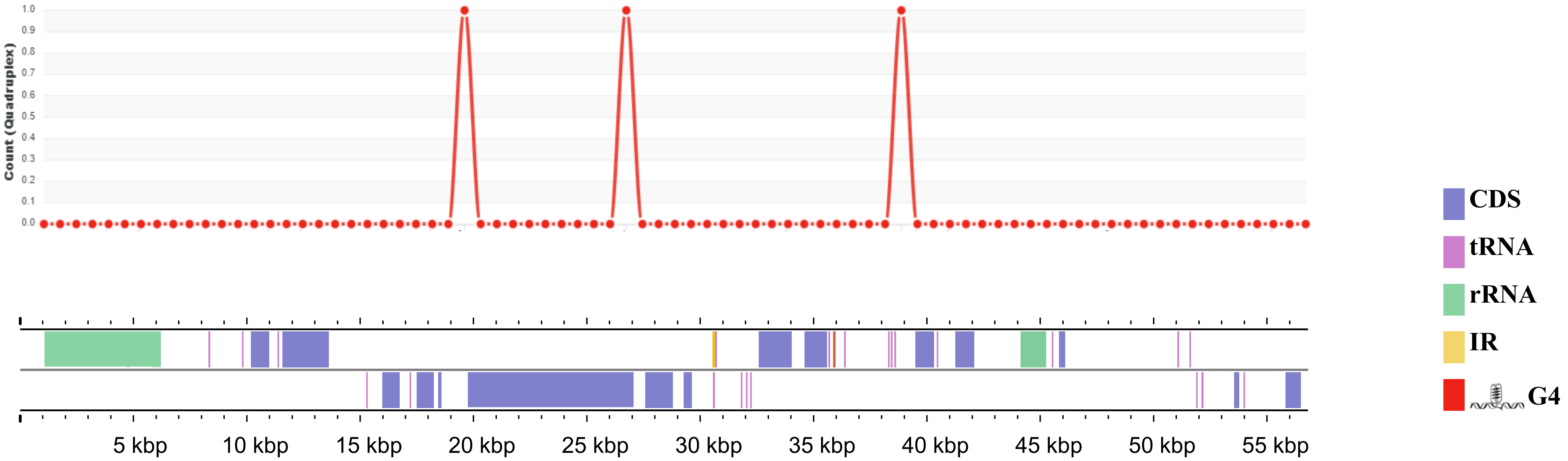
